# Harnessing Nature’s nanoSecrets: biocompatibility, biodistribution and bioactivity of extracellular vesicles derived from microalgae

**DOI:** 10.1101/2023.04.04.535547

**Authors:** Giorgia Adamo, Pamela Santonicola, Sabrina Picciotto, Paola Gargano, Aldo Nicosia, Valeria Longo, Noemi Aloi, Daniele P. Romancino, Angela Paterna, Estella Rao, Samuele Raccosta, Rosina Noto, Monica Salamone, Irene Deidda, Salvatore Costa, Caterina Di Sano, Giuseppina Zampi, Svenja Morsbach, Katharina Landfester, Paolo Colombo, Mingxing Wei, Paolo Bergese, Nicolas Touzet, Mauro Manno, Elia Di Schiavi, Antonella Bongiovanni

## Abstract

Nanoalgosomes are extracellular vesicles (EVs) released by microalgal cells that can mediate intercellular and cross-kingdom communication. In the present study, starting from the optimized nanoalgosome manufacturing from cultures of marine microalgae, we evaluated their innate biological properties in preclinical models. Our investigation of nanoalgosome biocompatibility included toxicological analyses, starting from studies on the invertebrate model organism *Caenorhabditis elegans,* proceeding to hematological and immunological evaluations in mice and immune-compatibility *ex vivo*. Nanoalgosome biodistribution was evaluated in mice with accurate space-time resolution, and in *C. elegans* at cellular and subcellular levels. Further examination highlighted the antioxidant and anti-inflammatory bioactivities of nanoalgosomes. This holistic approach to nanoalgosome functional characterization showcases that nanoalgosomes are innate effectors and potential drug delivery system for novel cosmetic formulations and EV-based therapies.

## INTRODUCTION

Cell-free therapy has emerged as a promising approach in the field of biomedical research, offering innovative strategies for the treatment of various diseases^1–3^. Among the major players in this field, extracellular vesicles (EVs) stand out as promising cell-derived therapeutic effectors^1,3–5^. Within naturally occurring nanoparticles, EVs have shown immense potential as intercellular mediators due to their ability to transfer bioactive cargo molecules to recipient cells and thereby influence their behaviour^1,3,6^. This cargo includes growth factors, cytokines, microRNAs and other signalling molecules that can modulate cellular processes and promote cell survival, proliferation, differentiation, tissue repair and regeneration^7^. The therapeutic potential of EVs is influenced by their cellular origin, resulting in different changes in the target cells. For example, EVs derived from human mesenchymal stem cells (MSCs) have shown regenerative properties, while EVs derived from cancer cells promote tumor progression^4,7^. Another important aspect to consider when using EVs as therapeutic effectors is their immunomodulatory properties^8–10^. Milk-derived EVs, for instance, have remarkable regenerative properties due to their high content of growth factors and immunomodulatory molecules^9^. Plant-derived EVs also show therapeutic potential due to their bioactive cargo, including phytochemicals and secondary metabolites that can influence immune responses and promote an anti-inflammatory environment^11,12^. In addition, EVs derived from microorganisms such as bacteria offer an interesting opportunity for therapeutic applications, particularly as vaccines, as they can carry a wide range of functional molecules and have unique bioactivity derived from their microbial origin that promotes enhanced immune activation^13^. Furthermore, EVs can be loaded with exogenous therapeutic entities (such as small molecule drugs, nucleic acid, proteins, and CRISPR/Cas9) and their endogenous cargo can intensify the effect of the encapsulated drugs, creating a combinatorial effect^14–16^. In recent years, interest in EVs derived from human MSCs has increased due to their therapeutic potential in preclinical studies across regenerative medicine in diverse tissues, such as lung, kidney, liver, central nervous system, cartilage, bone, and heart^17^. However, the therapeutic potential of MSC-EVs is still debated due to the complexity of MSCs, including their tissue origin and cell culture conditions. As the production of MSC-EVs has proven to be neither sustainable nor controllable, as well as economically and environmentally viable, the academic and industrial communities have been investigating alternative approaches to obtain EVs from other biological sources^12,18–22^. In this context, we are exploring a fascinating advance in the field, showcasing a new specific type of EVs that we call “nanoalgosomes” or “algosomes”^23^. Nanoalgosomes are small EVs (sEVs) isolated from microalgae conditioned media, that are surrounded by a lipid bilayer membrane, contain EV biomarkers and have a typical EV size distribution, morphology and density, and are highly stable in human blood plasma, non-cytotoxic *in vitro* and can be taken up by various cellular systems^23,24^. We focused on small EVs isolated from the photosynthetic marine chlorophyte *Tetraselmis chuii*, which is rich in vitamin E, carotenoids and chlorophyll with anti-inflammatory and antioxidant properties^23–25^. Through our patented platform, we have achieved high-yield production of quality-controlled EVs from microalgae, a bioresource that can be grown in scalable, renewable and environmentally sustainable photo-bioreactors^26^. The current study aims to fully elucidate the biological properties of nanoalgosomes and to validate their safe use as innate bioactive effectors in EV-based therapy, through an original multi-branched approach starting with studies on living organisms and delving into molecular mechanisms at the subcellular level. Our analyses show that nanoalgosomes are biocompatible and exert anti-inflammatory and antioxidant effects both *in vitro* as well as *in vivo*, using mouse models and the invertebrate model organism *Caenorhabditis elegans*, a faster and less expensive alternative to mammalian models, whose body transparency is advantageous for studying nanoparticle toxicity, distribution and uptake in living organisms^25^. Understanding and exploiting the potential of nanoalgosomes as natural bio-based nanoparticles with innate bioactivity could pave the way for safe, innovative, and effective formulations that would benefit various fields of nanomedicine, as well as cosmetics.

## RESULTS AND DISCUSSION

### BIOLOGICAL FEATURES OF NANOALGOSOMES

#### Production and quality checking of nanoalgosomes

Firstly, we considered the guidelines of MISEV 2018 to quality check each nanoalgosome preparations used for subsequent biological evaluations^27^. Multiple orthogonal physical and molecular techniques have been applied, including Nanoparticle Tracking Analyses (NTA), fluorescent-NTA (F-NTA) following membrane-labeling of nanoalgosomes, Dynamic Light Scattering (DLS), cryo-Transmission Electron Microscopy (cryo-TEM), Atomic Force Microscopy (AFM), and immunoblotting analyses of nanoalgosome markers (Figure 1 a-f). This approach allowed to demonstrate the presence of small EVs with expected average diameter (e.g., 100±10 nm measured by NTA and DLS), morphology, EV markers (*e.g.*, Alix, H+-ATPase and Enolase positivity) and proper EV protein/particle number ratio (*e.g.*, 1 µg of total EV protein corresponds to a range of 5-10×10^9^ in all nanoalgosome batches, n=6)^28^. All these features grant batch-to-batch consistency and quality, and repeatedly yielding vesicles (∼10^12^ nanoalgosomes/L of microalgal conditioned-medium, corresponding to ∼10^4^ nanoalgosomes/microalgal cell) with stable biophysical and biochemical properties. These results on nanoalgosomes are consistent with previous studies, and emphasize the importance of accurate quality control checking of nanoalgosome preparations to further exploit their potential as innovative therapeutic agents^27,29^.

**Figure 1.**
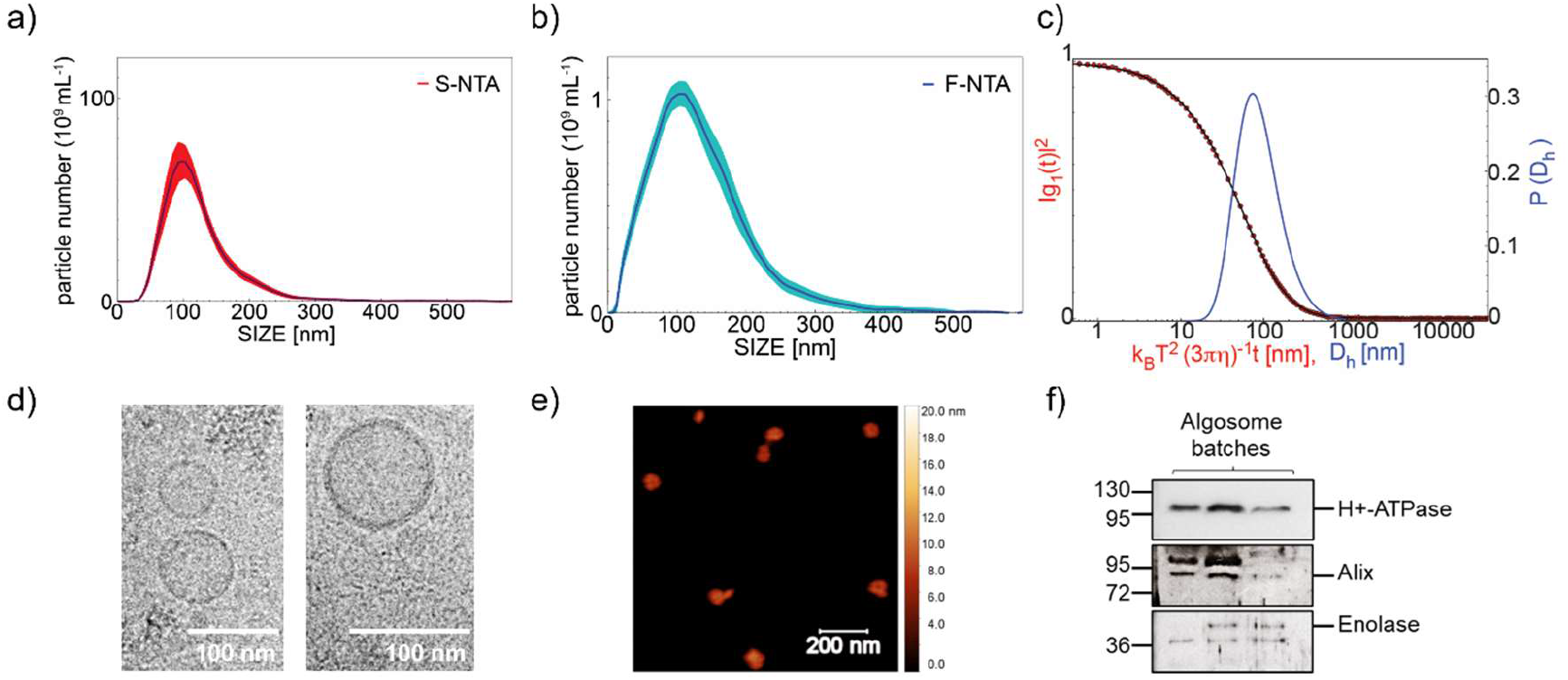
Quality control of nanoalgosome preparations. The particle size and concentration of the nanoalgosome preparations were determined by a) NTA, b) F-NTA and c) DLS. Morphology of nanoalgosomes was analyzed by d) cryo-TEM and e) AFM. Immunoblot analyses were performed on nanoalgosome batches to detect the markers H^+^-ATPase, Alix and Enolase. Representative results of three independent biological replicates are presented.

#### Nanoalgosome internalization mechanism in vitro and in vivo

In our previous study, we demonstrated that nanoalgosomes, when labeled with different fluorescent dyes (*i.e.*, Di-8-ANEPPS, PKH26, or DiR), are taken up and localized *in vitro* in the cytoplasm of cells and *in vivo* in the cytoplasm of *C. elegans* intestinal cells^24,25^. Therefore, we investigated the molecular mechanisms involved in nanoalgosome internalization *in vitro* and *in vivo*. Since we proved that nanoalgosomes are internalized within human cells in a dose- and time-dependent manner through an energy-dependent mechanism, we hypothesized active endocytic pathways, including macropinocytosis, clathrin- and caveolae-mediated endocytosis, as possible mechanisms of nanoalgosome internalization that were reported for other EVs^30^. To test among these alternative hypotheses, we used three specific blocking agents in 1-7 HB2 cells: dynasore to inhibit clathrin-mediated endocytosis, nystatin to interfere with caveolae-dependent uptake, and 5-[N-ethyl-N-isopropyl] amiloride (EIPA) to inhibit macropinocytosis^31–34^. We monitored intracellular nanoalgosome uptake in cells by measuring the fluorescence intensity of Di-8-ANEPPS-labeled nanoalgosomes after 2 and 3 hours of incubation, with or without different inhibitor treatments (Figure 2a, Supporting Figure 1a-b). The results showed that cells treated with dynasore (60µM) had a nanoalgosome internalization trend similar to the negative control (*i.e.,* cells incubated at 4°C, which are inhibited for all energy-dependent processes), thus indicating that dynasore inhibited nanoalgosome cellular uptake. In contrast, no significant endocytosis inhibition was observed in cells treated with EIPA (10µM) or nystatin (50µM), which showed a nanoalgosome internalization level similar to the positive control (*i.e.*, cells incubated at 37°C). Cells are viable for each treatment, demonstrating that none of the inhibitors used, at any concentration, were toxic to the cells (Supporting Figure 1a). These results indicate that clathrin-dependent endocytosis plays a role in the cellular uptake of nanoalgosomes.

**Figure 2.**
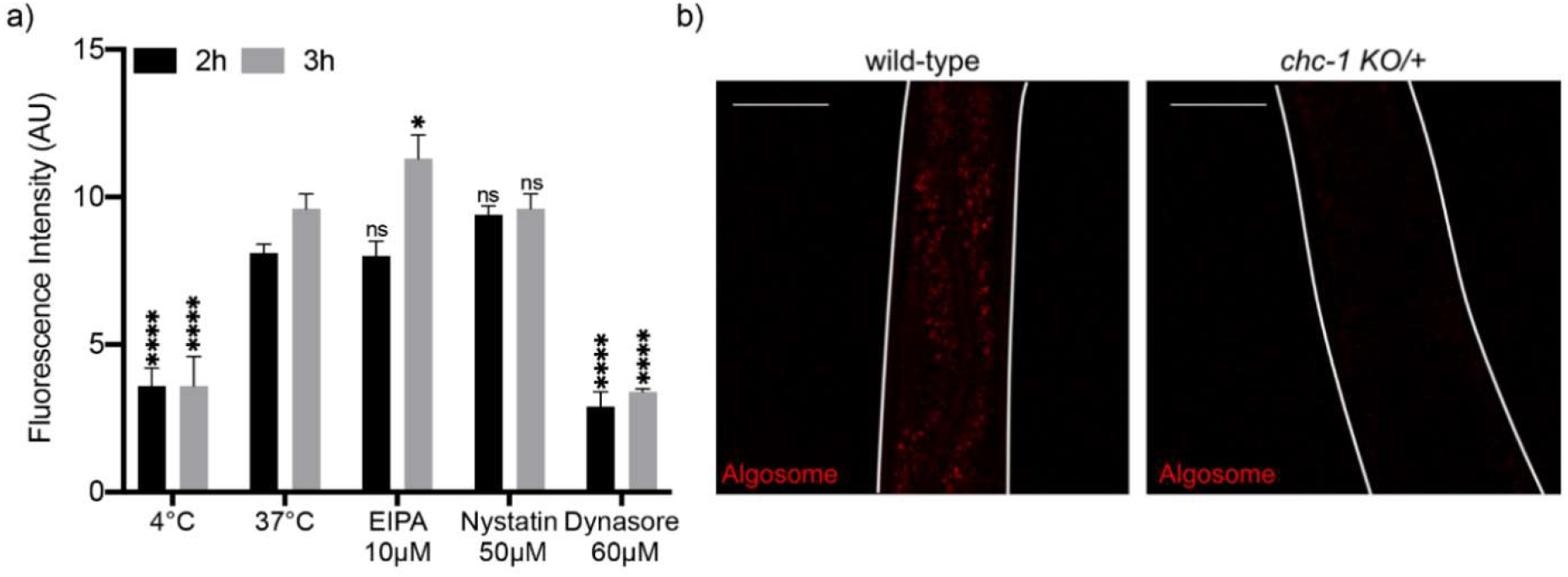
Molecular mechanism of nanoalgosome internalization in human cell lines and *in vivo*. a) Effect of metabolic inhibitors of endocytosis on nanoalgosome uptake in 1-7 HB2 cells treated with nystatin, EIPA and dynasore. Results are presented as arbitrary unit of nanoalgosome fluorescence intensity inside cells after 2 and 3h of incubation. Each value represents the mean ± SD from three independent experiments. One-way ANOVA statistical test was used to assess the statistical significance of the differences: 37°C (2h) (3h) vs 4°C (2h) (3h), EIPA 10 µM (2h) (3h), nystatin 50 µM (2h) (3h) and dynasore 60 µM (2h) (3h), ****p<0,0001, ns=not significant. b) Representative confocal images of wild-type (left), *chc-1 KO* (right) heterozygote animals after 24h of treatment with Di-8-ANEPPS labeled nanoalgosomes (red). Fluorescent signal is observed only in the intestinal cells of wild-type animals, while in *chc-1 KO* heterozygote animals the fluorescent signal is strongly reduced. Animal body is outlined with white lines. Scale bar is 50 μm.

To confirm the results obtained *in vitro,* we used the model system *C. elegans*, taking advantage of the ease to use genetic mutants and dissect molecular pathways in this model. In particular, when the clathrin heavy chain gene is mutated in *C. elegans*, the uptake of synthetic nanoparticles is impaired^35^.

Thus, we used a *KO* mutant in the clathrin heavy chain, *chc-1(ok2369*), and since *chc-1* is an essential gene and its depletion causes animal lethality, we analyzed heterozygous balanced animals (*chc-1 KO/+*). We treated for 24 hours animals with nanoalgosome fluorescently labeled with Di-8-ANEPPS and observed, in wild-type animals, a fluorescent signal in the intestinal cells (Figure 2b, left panel). On the contrary, in all *chc-1 KO/+* animals analyzed, the fluorescent signal in the intestinal cells was strongly reduced (Figure 2b, right panel). The identification of clathrin-mediated endocytosis as the primary mechanism responsible for the cellular uptake of nanoalgosomes is aligned to previous studies, which investigated the routes and mechanisms of mammalian cell-derived EV internalization, highlighting the conservation of this endocytic mechanism towards EVs from different species^3,30^. The clathrin-mediated endocytic relies on receptor-mediated, hydro-phobic or electrostatic interactions in areas of clathrin expression on the cell membrane, and is also the most common route for synthetic nanoparticles and virus uptake in non-specialized mammalian cells^36,37^.

#### Nanoalgosome intracellular localization in vitro and in vivo

We further investigated the subcellular fate of labeled nanoalgosomes once internalized, both *in vitro* and *in vivo*. The subcellular localization of PKH26-labeled nanoalgosomes in MDA-MB 231 cell line was determined using immunofluorescence and confocal microscopy (Figure 3a). Three distinct subcellular compartments were evaluated for *in vitro* study: the endosomal compartment, the lysosomal system, and the endoplasmic reticulum (ER), using established biomarkers (CD63, LAMP1, and Calnexin, respectively).

**Figure 3.**
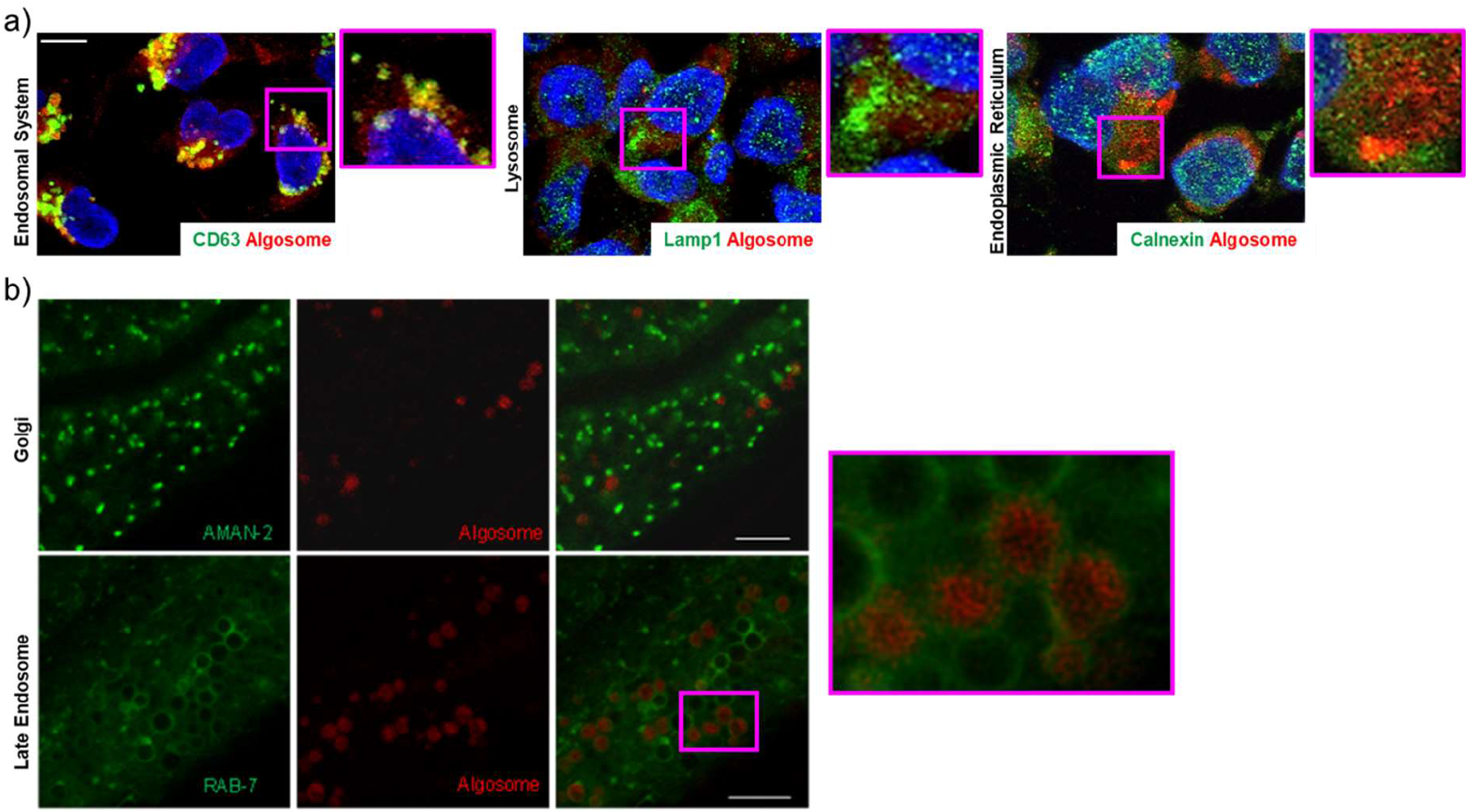
Intracellular nanoalgosome localization. a) Confocal microscopy analyses in MDA-MB 231 cells. In green are the endosomal system (CD63), the endoplasmic reticulum (calnexin) and lysosomes (Lamp1) of MDA-MB 231 cells, incubated with PKH26-labeled nanoalgosomes (red) for 24h. Scale Bar are 50µm. b) Confocal microscopy analysis of *C. elegans* intestinal cells. In green the Golgi marker AMAN-2 (upper panels) and late endosomes marker RAB-7 (lower panels). In red Di-8-ANEPPs labeled nanoalgosomes. Scale bar 25 µm. An enlargement of the pink box is visible on the right.

After 24 hours of incubation, we found a co-localization of nanoalgosomes (red signal) with the endosomal protein CD63 (green signal). Fluorescence images (Supporting Figure 2) showed this co-localization in a large portion of the cells, suggesting involvement in endosomal compartments. Confocal images (Figure 3a) further confirmed this direct relation with the endosomal system in the area where the fluorescence appears yellow.

Furthermore, the localization of PKH26-labeled nanoalgosomes was evaluated in relation to two other intracellular markers, LAMP-1 and calnexin (Figure 3a). Both fluorescence and confocal images showed that intracellular LAMP-1- and calnexin-positive compartments (green) did not co-localize with internalized PKH26-labeled vesicles, suggesting that lysosomes and ER were not involved in their intracellular trafficking.

To confirm these observations *in vivo*, we took advantage of *C. elegans*’ transparency and transgenics expressing fluorescent proteins in the Golgi (GFP fused to AMAN-2 protein) or in late endosomes (GFP fused to RAB-7 protein). AMAN-2 is a membrane protein of the Golgi, and we did not observe any co-localization with Di-8-ANEPPS-labeled nanoalgosomes, as shown in Figure 3b (upper panels). On the other hand, when we used a marker for late endosome membranes (RAB-7), we observed the nanoalgosome fluorescent signal inside the endosomes (Figure 3b, lower panels). Taken together, our results show that after nanoalgosomes are internalized by clathrin-mediated endocytosis, the fluorescent signal of nanoalgosome membrane co-localizes only with the endosome and not with the lysosomes, thus demonstrating that nanoalgosomes are not destined to lysosomal degradation.

### NANOALGOSOME BIOCOMPATIBILITY *IN VITRO,* AND *IN VIVO*

#### In vitro *and* in vivo *genotoxicity*

Our group recently demonstrated that nanoalgosomes did not elicit cytotoxicity, hepatotoxicity or genotoxicity in different cell lines^23,24^. Here, we have examined whether nanoalgosome treatment could trigger the activation of DNA-damage pathways by conducting gene expression analysis in 1-7 HB2 normal mammary epithelial cells. Typically, DNA damage leads to cell cycle arrest, regulation of DNA replication, and activation of the repair pathway. DNA damage triggers the activation of DNA-damage sensors such as ATR (ATM and Rad3-related serine/threonine kinase) and their recruitment to DNA damage sites^38,39^. In addition, checkpoint kinase 1 (CHEK1) is a key downstream molecule of DNA-damage response signaling; CHEK1 phosphorylates various intracellular substrate proteins, including the RAD51 recombinase which is central to the homologous recombination pathway, and binds single-stranded DNA at damage sites, forming filaments observed microscopically as nuclear foci. Therefore, these genes are considered to be involved in the response to DNA damage maintaining genome integrity, with ATR initiating a signaling cascade that activates *CHEK1* and *RAD51*^40^. Gene expression analysis of 1-7 HB2 cells treated with 2 µg/mL of nanoalgosomes for 24 hours showed not significant changes in the expression level of these selected DNA-damage sensors (*ATR, CHECK1, RAD51*), thus suggesting that the outcomes related to nanoalgosomes are negligible on the activation of the DNA damage signaling pathway (Figure 4a).

**Figure 4.**
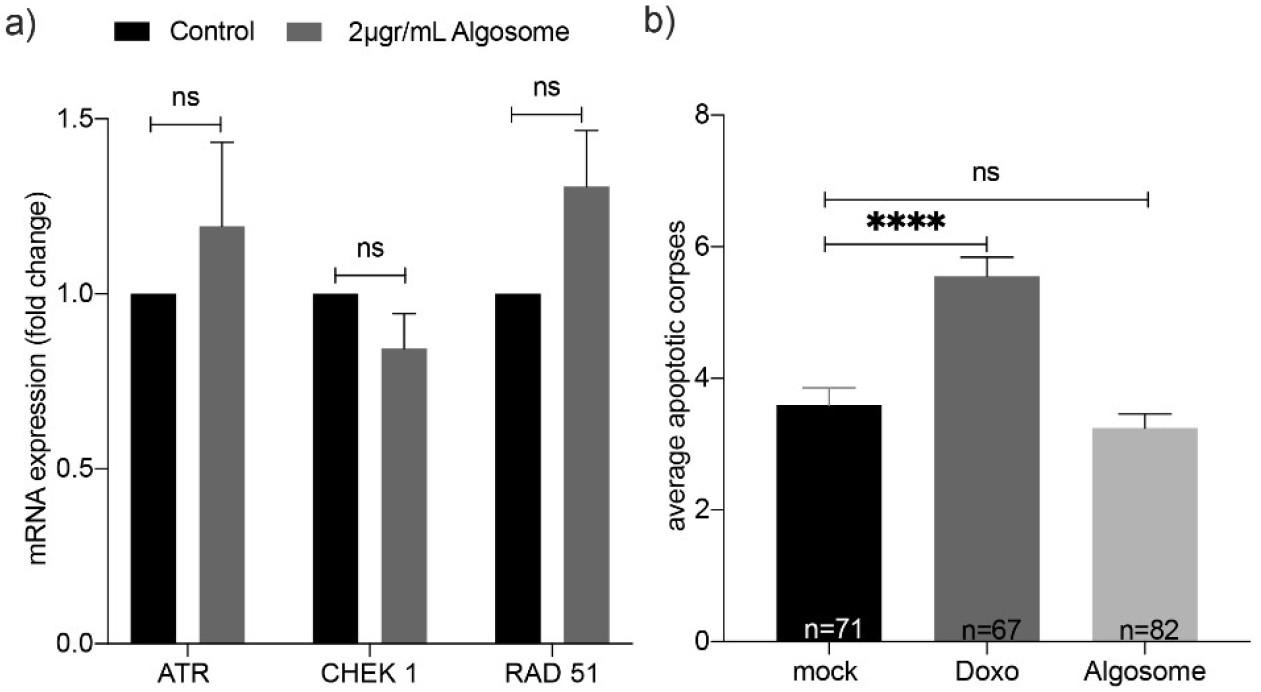
Genotoxicity assessment *in vitro* and *in vivo*. a) Real-time PCR for quantification of DNA-damage sensors expression levels following exposure to nanoalgosomes (2 μg/mL for 24h in 1-7 HB2 cells). Fold-changes were normalized with β-actin expression and given as relative to control. Each experiment was performed in triplicate. By One-way ANOVA differences of treated cells were determined not statistically significant when compared with the control (p>0.05). b) Quantification of SYTO12 positive apoptotic corpses in the germline after treatment with mock (PBS), Doxo (doxorubicin 2.4 μM), and nanoalgosomes (20 μg/mL). Bars represent means; error bar is the SEM. n is the number of animals analyzed. Kruskal-Wallis one-way ANOVA was used to assess the statistical significance of the differences: mock vs nanoalgosomes ns, p=0.7870; mock vs Doxo****, p<0.0001.

For *in vivo* analyses of nanoalgosome genotoxicity, we used *C. elegans* that allows the evaluation of the putative toxic effect of nanoparticles on a whole living animal, with fewer ethical concerns, lower costs, and an important reduction in the number of vertebrate animals used. Specifically, the *C. elegans* germline can be used as a tool to study genotoxicity *in vivo*^41^. Physiological germline apoptosis occurs in wild-type animals in absence of any stress and can be visualized as an average of three apoptotic corpses per gonadal arm using SYTO12, an apoptotic-DNA fluorescent marker^42,43^. Genotoxic agents, such as doxorubicin, are capable of increasing germline apoptosis as consequence of DNA damage^44^.Thus, after chronic treatment of 72 h of animals with nanoalgosomes at 20 µg/mL, we evaluated the germline apoptosis to assess nanoalgosomes genotoxicity *in vivo*. We did not observe any increase in the number of apoptotic corpses in the germline compared to mock (Figure 4b). Differently, animals treated with doxorubicin showed a higher number of apoptotic corpses compared to animals treated with mock, thus confirming that nanoalgosomes have no genotoxic effect at the concentration tested neither *in vitro* nor *in vivo*.

#### In vivo biocompatibility of nanoalgosomes in C. elegans

*C. elegans* offers a comprehensive set of experiments that allow to address the nanoparticle biocompatibility at different levels checking their effect on animal survival, growth, lifespan, fertility, lethality, and neuron viability, among others. Moreover, the short life cycle (three days) and lifespan (three weeks) of this model permits to exploit all these phenotypes in a very short period of time on large number of animals. Thus, we treated wild-type animals through *in liquido* culturing for 72 hours with increasing concentrations of nanoalgosomes (1, 20, 64, 128 µg/mL), first evaluating animal viability and capability of reaching adulthood. After treatment for three days *in liquido*, we did not observe any toxic effect of nanoalgosomes, as all the animals were alive and reached adulthood, similarly to the animals treated with PBS as mock (Figure 5a-b). Thereafter, we decided to use the concentration of 20 µg/mL to further evaluate nanoalgosome effects on animal fitness^45^.

**Figure 5.**
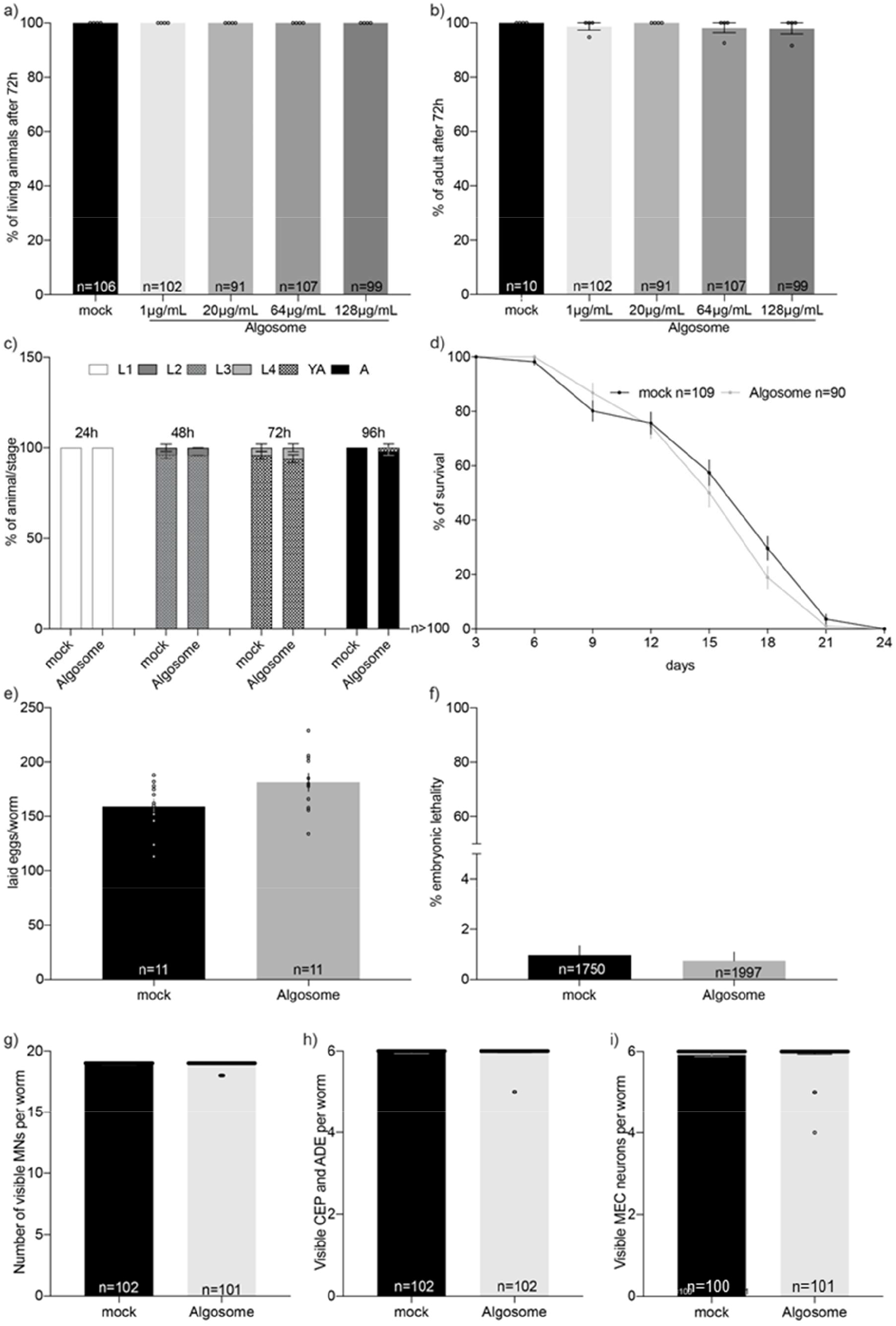
Nanoalgosome *in vivo* biocompatibility in *C. elegans*. a) Quantification of the number of living animals after treatment with mock (PBS) or different nanoalgosome concentrations. b) Quantification of the number of animals reaching adult stage after treatment with mock or different nanoalgosome concentrations. c) Quantification of the percentages of animals at each developmental stage, which are larval (L1-L4) and adult (Young Adult, YA, and Adult, A) stages, after treatment with mock and nanoalgosomes (20 µg/mL). d) Lifespan of animals treated with mock and nanoalgosomes (20 µg/mL). e) Brood size of animals after treatment with mock and nanoalgosomes (20 µg/mL). f) Embryonic lethality of animals after treatment with mock and nanoalgosomes (20 µg/mL). g-i) Quantification of the number of visible motoneurons (MNs), dopaminergic neurons in the head (CEP and ADE) and mechanosensory neurons (MEC) after treatment with mock and nanoalgosomes (20 µg/mL). Bars represent the means and dots replicates (in a-d, f-i) or single animals (in e); error bars are SEM. n is the number of animals analyzed (in a-d, i), the number of P0 animals (in e), or the number of eggs analyzed (f). In all graphs the statistical significance of the differences between treatments with mock and nanoalgosomes was assessed with Kruskal-Wallis One-way ANOVA (a-b), non-parametric Compare two proportions test (c), Mann Whitney *t-test* (e-i) or Log-rank (Mantel-Cox) test (d) and never found significant (p>0.05).

*C. elegans*’ development and growth are finely regulated in four larval stages followed by a fertile adult stage, which can be altered when the animals are exposed to toxic agents^46^. To evaluate animal growth, we treated animals chronically with nanoalgosomes and tested the percentage of animals at each developmental stage every 24 hours, without observing any alterations or delays compared to animals treated with mock (Figure 5c). We treated animals over their entire lifespan and, also in this case, nanoalgosomes had no effect on their survival curve (Figure 5d).

Another aspect used to assess the biocompatibility of nanoparticles is to evaluate their effects on animal fertility and offspring survival. Therefore, after chronic treatment of animals from eggs to L4 larval stage, we assessed the animals’ egg-laying ability and egg hatching (Figure 5e-f), without observing any effects of nanoalgosome treatment compared to mock. This result confirms that nanoalgosomes do not affect animal fertility, offspring release, or embryonic survival. Moreover, we decided to evaluate the putative neurotoxicity of nanoalgosomes. *C. elegans*’ transparency makes it possible to observe specific cell types, such as neurons, in living animals by using fluorescent proteins that are only expressed in the cells of interest. We thus evaluated the effects of nanoalgosomes on three different neuronal classes: GABAergic motoneurons, dopaminergic sensory neurons, and glutamatergic mechanosensory neurons. Interestingly, the invariant cell lineage in *C. elegans* determines a fixed number of neurons in adults, with 19 GABAergic motoneurons in the ventral cord, 6 dopaminergic neurons in the head, and 6 mechanosensory neurons along the body. Chronic treatment with nanoalgosomes did not cause any death of the neurons analyzed (Figure 5g-i; Supporting Figure 3a-c) as the number of visible neurons was not affected by the treatment. Moreover, all these neurons exhibited normal morphology.

Taken together, our results demonstrate that nanoalgosomes are very well tolerated *in vivo* by *C. elegans*, and no toxicity was observed after chronic exposure from eggs to adult stage and throughout animal life at these concentrations.

#### In vivo biocompatibility of nanoalgosomes in wild-type mice

Biochemical and hematological studies were conducted on wild-type mice to thoroughly evaluate the *in vivo* biocompatibility in immune-competent mice, after a single intravenous (I.V.) administration of nanoalgosomes. Wild-type BALB/c mice (n=4, 2 male and 2 female) were injected with nanoalgosomes and a negative control (i.e., PBS), via intravenous injection into the retroorbital sinus. The starting nanoalgosome doses were based on literature; in different pre-clinical studies, the dose of EVs per kg of body weight ranged from 0.10 to 100 mg of total EV proteins, with an average of 2.75 mg/kg^47,48^. Based on the estimation that the body weight of 6-week-old BALB/c mouse is approximately 20 grams, the average I.V. dose of EVs in mice corresponds to 55 µg/mouse^49,50^. Thus, in our pre-clinical studies we used the following doses: Dose 1 (low dose) = 10 µg/mouse, corresponding to 4×10^10^ EVs/mouse; Dose 2 (high dose) = 50 µg/mouse, corresponding to 2×10^11^ EVs/mouse. Blood samples were collected from mice to analyze hematocrit, creatinine, blood urea nitrogen (BUN), liver transaminase enzymes (Serum Aspartate Transaminase, AST, and Serum Alanine Transaminase, ALT), and lymphocyte numbers. Table 1 shows slight changes in white and red blood cell counts, hemoglobin, or hematocrit in nanoalgosome-treated mice compared to control mice (*i.e.*, injected with PBS) at 48 hours post-IV injection; these differences are still within the normal range of values reported for BALB/c mice (Janvier Labs, France), thus suggesting the absence of potential immune-reaction to nanoalgosomes.

**Table 1.** Hematological and biochemical analyses of peripheral blood from BALB/c mice.

| Parameters | Dose 1 | Dose 2 | Control |
| --- | --- | --- | --- |

| Hematological |  |  |  |
| --- | --- | --- | --- |
| Red blood cells (x10<sup>6</sup>/μL) | 10.1 ± 0.0 | 9.9 ± 1.0 | 10.1 ± 0.1 |
| Hemoglobin (g/dL) | 15.2 ± 0.0 | 15.0 ± 1.4 | 15.5 ± 0.5 |
| MCV (fL) | 47.0 ± 0.2 | 45.5 ± 0.5 | 47.5 ± 0.5 |
| Hematocrit (%) | 47.0 ± 0.2 | 46.0 ± 5.0 | 48.0 ± 1.0 |
| Reticulocytes (%) | 3.4 ± 0.1 | 2.1 ± 0.0 | 2.2 ± 0.3 |
| MCH (pg) | 15.0 ± 0.0 | 15.0 ± 0.0 | 15.5 ± 0.5 |
| Leukocytes (x10<sup>3</sup>/μL) | 3.1 ± 0.0 | 3.1 ± 2.0 | 4.8 ± 0.9 |
| Lymphocytes (%) | 72.0 ± 0.3 | 67.0 ± 2.0 | 78.0 ± 2.0 |
| Neutrophils (%) | 20.0 ± 0.1 | 25.0 ± 1.0 | 16.0 ± 2.0 |
| Monocytes (%) | 2.0 ± 0.0 | 3.0 ± 1.0 | 1.5 ± 0.5 |
| Eosinophils (%) | 3.0 ± 0.0 | 3.4 ± 0.5 | 2.5 ± 0.5 |
| Basophils (%) | 0.0 ± 0.0 | 0.5 ± 0.0 | 0.5 ± 0.0 |
| Biochemical |  |  |  |
| Creatinine (mg/L) | 4.0 ± 0.0 | 4.0 ± 0.8 | 4.2 ± 0.4 |
| BUN (mg/dL) | 23.0 ± 5.7 | 20.6 ± 6.6 | 27.0 ± 2.2 |
| Urea (g/L) | 0.5 ± 0.1 | 0.4 ± 0.1 | 0.5 ± 0.0 |
| ALT (x10<sup>3</sup> UI/L) | 2.8 ± 1.5 | 2.6 ± 0.4 | 2.1 ± 0.4 |
| AST (x10<sup>3</sup>UI/L) | 2.5 ± 0.6 | 4.0 ± 0.6 | 3.4 ± 0.3 |

This finding was corroborated by the *ex vivo* basophil activation test using whole blood from human healthy subjects (n=3), which assesses the activation state of these cells with or without stimulation^51^. In this test, a specific monoclonal antibody (anti-FcɛRI), binding to the high affinity IgE receptor, was used as positive control. Flow cytometry characterization showed no basophil activation following nanoalgosome treatments at different doses (0.25-2 µg/mL), with a percentage of basophil activation similar to the negative control (Supporting Figure 4; Supporting Table 1), thus demonstrating the absence of acute immediate allergic reactions induced by nanoalgosomes.

Further, creatinine, urea and BUN, as well as AST and ALT values were similar to the control group, suggesting that the nanoalgosome treatment does not impair normal kidney and liver functions. These results were paralleled with the lack of any obvious histological changes in the liver and spleen architecture after 48 hours of nanoalgosome I.V. administration (Supporting Figure 5). Thus, both two doses of nanoalgosomes did not exhibit toxic effects, blood parameter alterations after 48 hours following a single acute I.V. administration.

The effects of nanoalgosomes in mice were also evaluated through clinical signs, body weight, and visual observations. Daily clinical examinations of treated mice were performed to exclude any behavior and signs of suffering, such as cachexia, weakness, difficulty in moving or feeding, hunching and convulsions. Further, supporting Table 2 shows that there was not significant body weight loss 3 and 6 days following nanoalgosome injection., indicating that nanoalgosome administration was well-tolerated by the animals. Our results demonstrate that a single dose of up to 2×10^11^ nanoalgosomes per mouse did not elicit any noticeable local and systemic toxicity in immune-competent BALB/c mice, allowing for dose escalation or repeated administrations. This is in line with our *C. elegans* results and with previously published data on I.V. administration in mice of xenogeneic milk-, human- and plant-derived EVs^8,10,11^. However, additional safety evaluations are necessary, since it is important to further expand *in vivo* studies on nanoalgosome tolerability with repeated administration in mice and larger animals, along with biodistribution information to fully understand the potential and future applications^48^.

### BIODISTRIBUTION OF NANOALGOSOMES IN NUDE MICE

Due to the presence of a lipid bilayer membrane, EVs can be labeled with fluorescent lipid dyes and their biodistribution has been evaluated in pre-clinical studies mainly using mouse models and recently using larger animals, including the pig-tailed macaque (*Macaca nemestrina*)^48,52,53^. To assess the biodistribution of nanoalgosomes systemically delivered in mice, the near-infrared dye DiR was used for nanoalgosome labeling. DiR is a lipophilic dye that fluoresces intensely only when inserted into a lipid membrane. The fluorescence spectrum of this dye (emission peak of 790 nm) allows for efficient penetration through bones and tissues with low autofluorescence, making it ideal for imaging in living animals. Nanoalgosomes were labeled with the DiR probe, while the same amount of DiR diluted in PBS was used as a free dye control. Both samples underwent NTA analysis after the removal of the free dye by ultracentrifugation and extensive washing steps. The NTA data showed a typical nanoalgosome size distribution with a mode size of 100 nm in diameter, after DiR labeling (Supporting Figure 6a). The presence of DiR-fluorescence in the nanoalgosome-labeled samples was verified by IR measurements using an Odyssey scanner, while for the free dye control, no fluorescence signal was detected (Supporting Figure 6b). Athymic nude mice (n=6; 3 male, 3 female) were used for these IVIS Spectrum Imaging studies, to minimize the possible fluorescence background and as they are largely used for xenogeneic EV administration; these mice were intravenous injected with aforementioned DiR-labeled nanoalgosomes at two doses (or the same volume of the free dye control)^52,54^. The biodistribution of the DiR-labeled nanoalgosomes was examined in live animals using IVIS. Prone and supine mice (prone male in Figure 6a-c; prone female and supine male in Supporting Figure 7a-b) were imaged at 3, 6, 24 and 48 hours post-IV injection. At 3 hours post-injection, the DiR-nanoalgosome fluorescent signals were accumulating mainly in the liver (as commonly observed for other EVs). Fluorescent signals increased in a time- and dose-dependent manner, as shown in the total radiant efficiency plot (Figure 6d). In contrast, lower and more constant fluorescent background signals were detected for the free dye control-treated mice after each imaging, highlighting the specificity of the DiR-nanoalgosome signal. Unexpectedly, nanoalgosomes were mainly localized in the bones 24 hours post-IV injection and, as shown in Figure 6e and f, their concentration significantly increased and significantly accumulated in femur by 48 hours. This organotropism is peculiar for nanoalgosomes, as most mouse studies with MSC-derived EVs have reported that EVs accumulate to the liver, spleen, and sometimes kidney and lung, with rapid clearance from blood circulation after 24h systemic injection^47,52,53^. Our data suggest a specific organotropism for bone compartments. Furthermore, the increasing fluorescent signals for up to 48 hours in mice and in individual areas (*i.e.*, backbone and femur) suggests that nanoalgosomes are stable in body fluids and that their circulating half-life is high.

**Figure 6.**
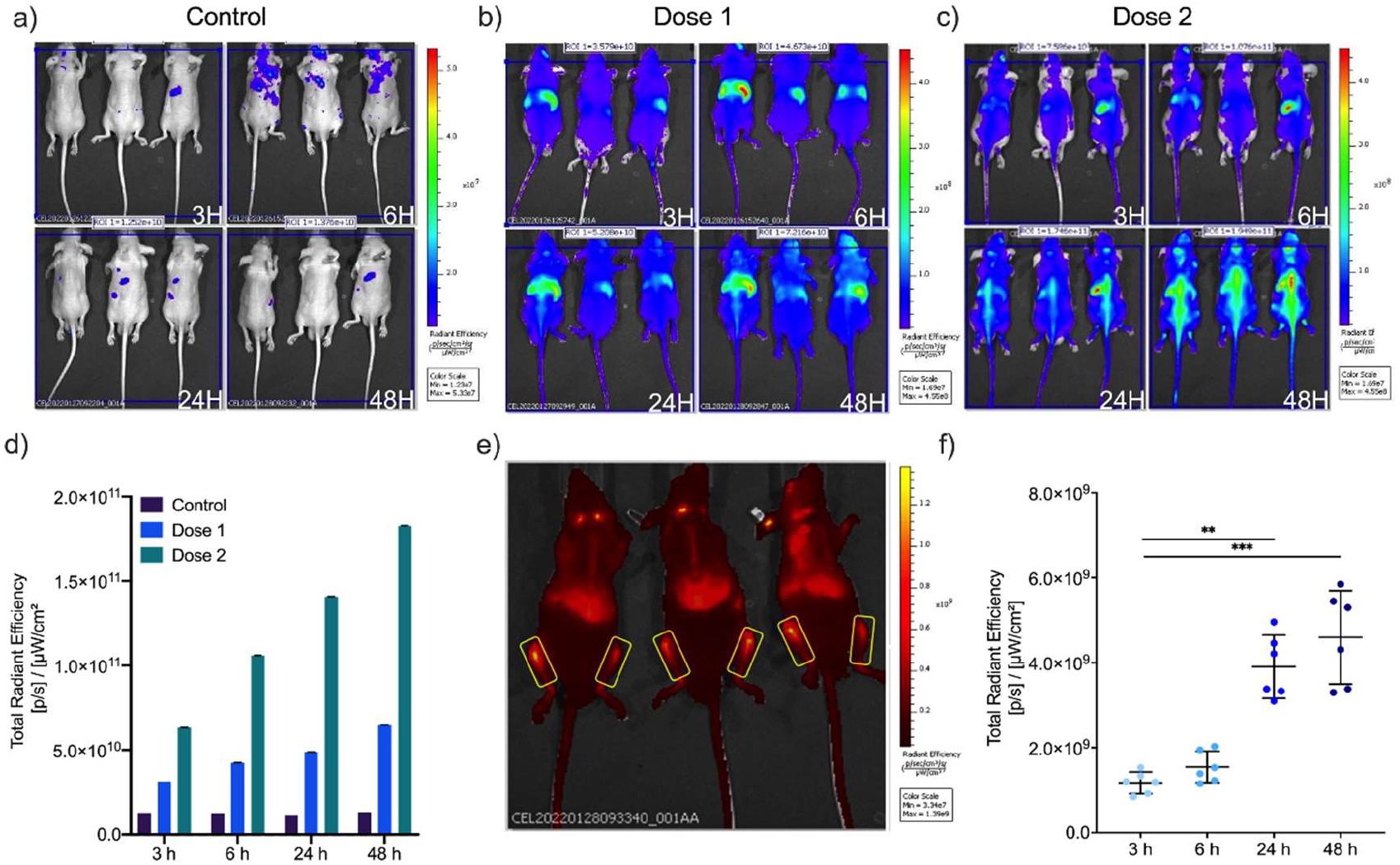
IVIS imaging of DiR-labeled nanoalgosome biodistribution in nude mice after IV injection. Nude mice (n=3; male, prone) were injected with (a) v/v of free dye control /200 µL/mouse; (b) dose 1: 10µg nanoalgosomes/200 µL/mouse; (c) dose 2: 50µg nanoalgosomes/200 µL/mouse and control. Nanoalgosome biodistribution was measured at 3, 6, 24 and 48h post-injection using an IVIS *in vivo* imaging system at ex/em 644/665. d) Total radiant efficiency plot of DiR-labeled nanoalgosomes (doses 1 and 2) and the free dye control 3, 6, 24 and 48h post-injection. e-f) Representative images of DiR-nanoalgososome fluorescence in the femur (48h post I.V.) and respective radiance efficiency quantification graph at 3, 6, 24 and 48h. Kruskal-Wallis one-way ANOVA was used to assess the statistical significance of the differences: 3h vs 24h**, p<0.01; 3h vs 48h***, p<0.001.

### NANOALGOSOME BIOACTIVITY *IN VITRO* AND *IN VIVO*

#### Anti-inflammatory activity of nanoalgosomes

The bioactivity of nanoalgosomes was investigated in immune-responsive macrophage cells. Initially, we checked the viability of THP-1 cells treated with different concentrations of nanoalgosomes for up to 72 hours, confirming that nanoalgosomes (up to 2 µg/mL) did not induce any cell toxicity (Supporting Figure 8). Subsequently, THP-1 cells were pre-treated with 0.5 µg/mL of nanoalgosomes for 4 hours and afterwards exposed to lipopolysaccharide (LPS)-induced inflammation for 20 hours. We first excluded a cytotoxic effect induced by these experimental conditions (Figure 7a), then we monitored interleukin-6 (IL-6), which is a marker of inflammation, using qRT-PCR and ELISA tests. The results in Figure 7b and c show no significant differences in IL-6 induction following 24 hours of nanoalgosome treatment compared to untreated cells; this result is in line with the *in vivo* data, previously shown, and is indicative of the immune-compatibility of nanoalgosomes. Further, figure 7b and c shows that nanoalgosomes significantly reduced IL-6 induction in LPS-treated THP-1 cells, leading to a 4.5-fold reduction in mRNA level and a 7-fold reduction in IL-6 production, and indicating their anti-inflammatory activity *in vitro*.

**Figure 7.**
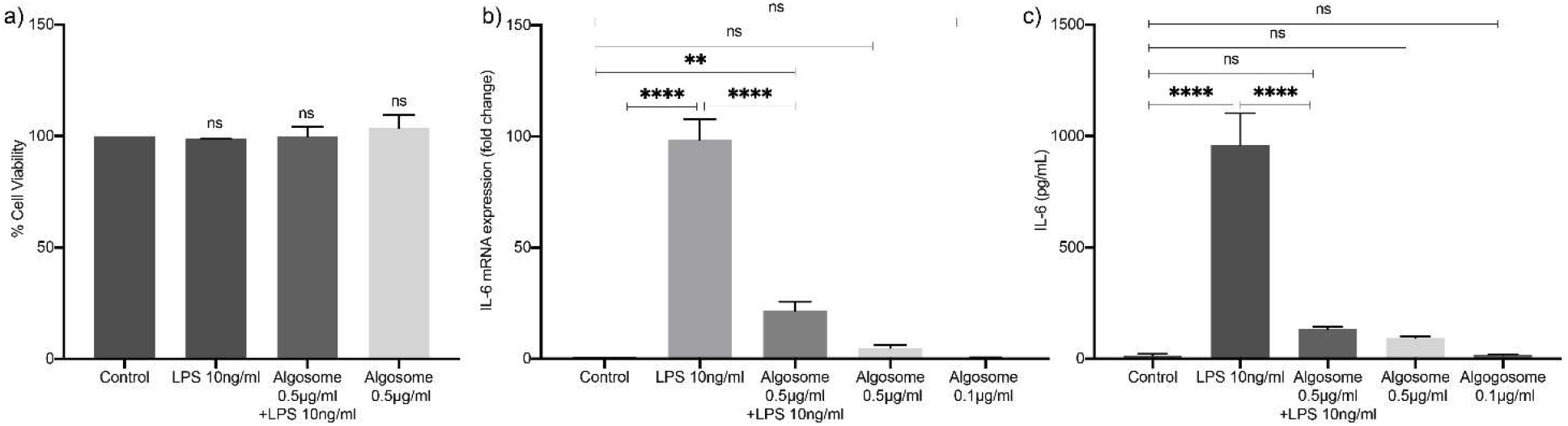
Anti-inflammatory effect of nanoalgosomes in immune-responsive cells. a) Cell Viability after exposure with LPS (10 ng/mL) and nanoalgosomes (0.5 µg/mL) for 24h, in TPH-1 cells. By Student’s t-test, differences of treated cells were not statistically significant when compared with the control. b) Real-time PCR quantification of *IL-6* mRNA relative levels after exposure with LPS (10 ng/mL) and nanoalgosomes (0.5 µg/mL) for 24h, in TPH-1 cells. c) ELISA results of IL-6 induction after exposure with LPS (10 ng/mL) and nanoalgosomes (0.5 µg/mL) for 24h, in TPH-1 cells. Two-way ANOVA statistical test was used to assess the statistical significance of the differences: Control vs LPS (10 ng/mL), LPS (10 ng/mL) vs Algosomes (0.5 µg/mL) + LPS (10 ng/mL), **p<0.01,****p<0.0001, ns=not significant. Representative results of three independent biological replicates.

#### Antioxidant activity of nanoalgosomes

Microalgae are sources of antioxidant compounds so we sought to determine the potential antioxidant role of microalgae-derived EVs^55,56^. Nanoalgosome antioxidant activity was evaluated in two different cell lines: tumoral (MDA-MB 231) and normal (1-7 HB2), using the cell-permeable 2′, 7′- dichlorofluorescein diacetate (DCF-DA) fluorescent probe that emits fluorescence proportionally to intracellular ROS content. Cell viability analyses showed that none of the concentrations of the oxidant agents used (i.e., H_2_O_2_ and tert-butylhydroperoxide, TBH) were toxic to the cells (Supporting Figure 9b-c). Figure 8a shows the percentage of ROS levels in cells treated with different nanoalgosome concentrations (0.5, 1, and 2 μg/mL) for 24 hours, with and without oxidative stress induction. As shown, treatments with nanoalgosomes per se did not induce oxidative stress, and the percentage of ROS levels was comparable to the negative control (untreated cells). After treatment with oxidant agents, ROS levels significantly increased in both cell lines. However, this increase in ROS levels was significantly lower in stressed cells which had been pre-treated with nanoalgosomes for 24 hours, suggesting that nanoalgosomes significantly reduced ROS levels and rebalanced the physiological ROS levels in both cell lines. To further investigate the antioxidant abilities of nanoalgosomes, we analyzed whether they could counteract ROS, directly or indirectly modulating the expression of oxidative stress responsive genes. We selected a panel of genes that play important roles in regulating oxidative stress and maintaining cellular homeostasis. For instance, AKR1C2 (aldo-keto reductase family 1 member C2) plays a role in detoxifying lipid peroxidation products, which can contribute to the production of ROS. FTH1 (ferritin heavy chain1) helps to regulate iron levels, which is important for ROS regulation since iron can catalyze the production of free radicals. Alox12 (arachidonate 12-Lipoxygenase) can contribute to oxidative stress by producing leukotrienes, which are inflammatory mediators that can increase ROS production. NOS2 (nitric oxide synthase) produces nitric oxide, which can have harmful effects on oxidative stress. Further, to counteract the effects of ROS, CAT, GPX1 and GSR (catalase, glutathione peroxidase and reductase, respectively) work together to neutralize ROS by converting them into less harmful products and to maintain the redox balance by reducing glutathione disulfide, which can be formed as a result of ROS exposure^57^. Thus, these proteins are interconnected and work together to regulate ROS levels in cells and maintain cellular homeostasis. After incubating 1-7 HB2 cells with 2 µg/mL of nanoalgosomes for 24 hours, with and without oxidant agent treatment, real-time PCR analyses were carried out to evaluate the mRNA expression levels of the selected genes involved in the oxidative stress cellular signaling (Figure 8b-d). The results showed that the expression of oxidative stress-related genes in cells treated with nanoalgosomes for 24 hours was similar to that of untreated cells, confirming that nanoalgosomes did not induce expression alterations of genes related to oxidative stress. As expected, after oxidative stress induction, these genes were upregulated, while interestingly the expression levels of most of the genes analyzed were significantly re-established or lowered in stressed cells pre-treated for 24 hours with nanoalgosomes. These results suggested that nanoalgosomes have potent antioxidant abilities, likely due to their antioxidant cargo and ability to neutralize free radicals, promoting protective mechanisms inside cells.

**Figure 8.**
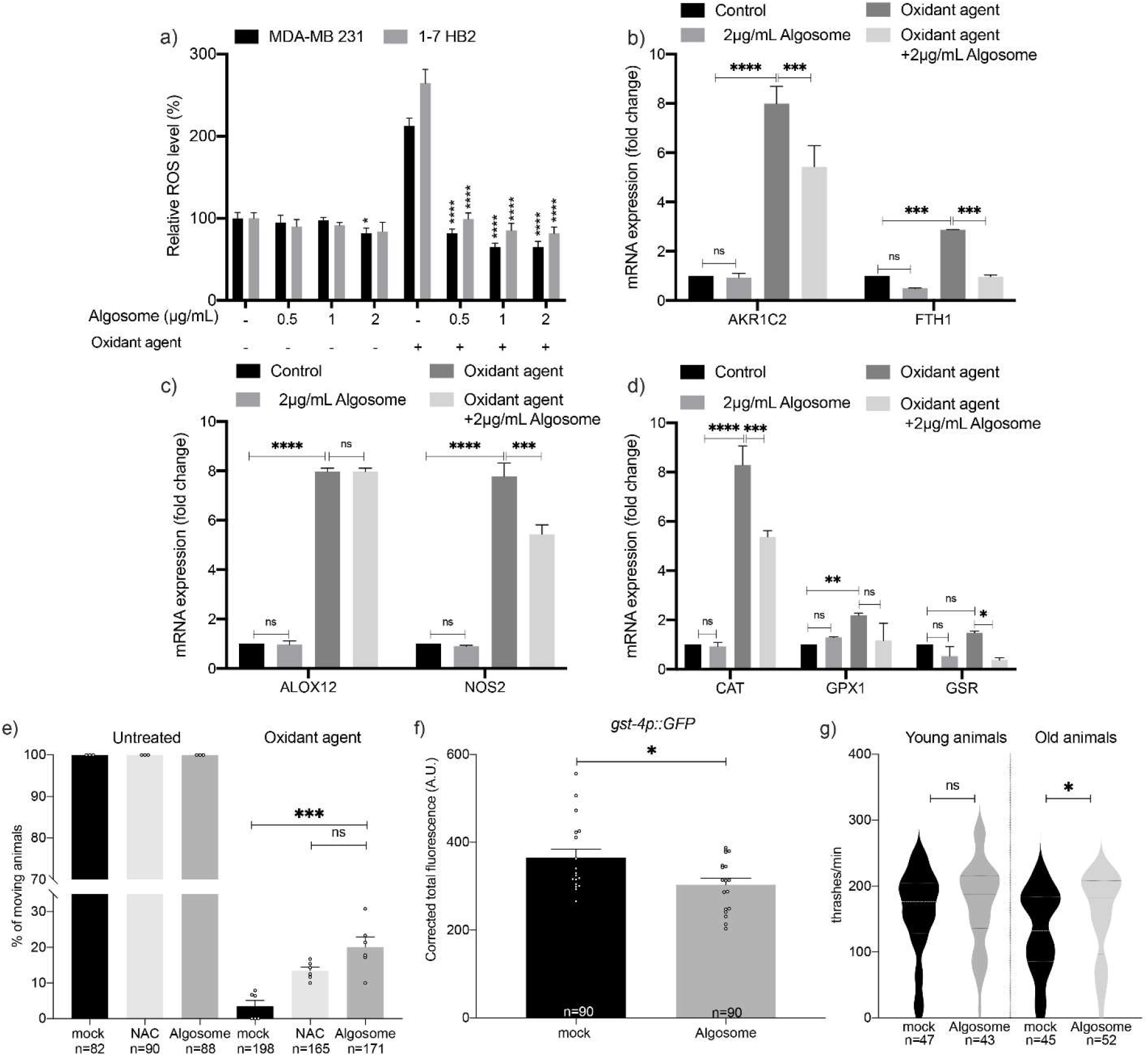
Antioxidant bioactivity of nanoalgosomes in cellular systems and in *C. elegans*. a) Percentage of ROS production in 1-7 HB2 and MDA-MB 231 cells treated with nanoalgosomes (0.5-1-2 µg/mL) for 24h, with/without oxidant agent (250 µM TBH), normalized to untreated cells. Values are means ± SEM of three independent experiments. One-way ANOVA was used to assess the statistical significance of the differences: MDA-MB 231 vs MDA-MB 231 2 µg/mL nanoalgosome, *p<0.05; 1-7HB2 vs 1-7HB2 2µg/mL nanoalgosome, *p<0.05; MDA-MB 231 + oxidant agent vs MDA-MB 231 0.5-1-2 µg/mL nanoalgosome + oxidant agent, ****p<0.0001; 1-7HB2 + oxidant agent vs 1-7HB2 0.5-1-2 µg/mL nanoalgosome + oxidant agent, ****p<0.0001. b-d) Real-time PCR quantification of antioxidant enzyme expression levels after exposure to nanoalgosomes (2 µg/mL) for 24h, with/without oxidant agent (250 µM H_2_O_2_) in 1-7 HB2 cells. Each experiment was performed in triplicate. Three-way ANOVA statistical test was used to assess the statistical significance of the differences: Control vs 2 µg/mL algosome; Control vs Oxidant agent; Oxidant agent vs Oxidant agent + 2 µg/mL algosome, *p<0.05, **p<0.01, ***p<0.001, ****p<0.0001, ns=not significant. e) *In vivo* response to exogenous oxidative stress. Animals have been treated with mock, NAC (2mM) and nanoalgosome (20 µg/mL). Animal movement was assessed right after acute exposure with water (untreated) or H_2_O_2_ 5mM. H_2_O_2_ mock vs H_2_O_2_ nanoalgosome ***p=0.0009; H_2_O_2_ nanoalgosome vs H_2_O_2_ NAC ns, p=0.5252. f) Quantification of the fluorescence in *gst-4p*::GFP expressing animals after chronic treatment with mock or nanoalgosomes (20 µg/mL). Each dot represents the total fluorescence in a picture corrected for the background. Bars represent the mean; error bar is the SEM. n is the total number of animals analyzed. *t-*test: mock vs nanoalgosome *p=0.0435. g) Thrashing assay on young and old animals after acute treatments with mock and nanoalgosomes (10 µg/mL). Violin plots show the distribution of thrashes performed by the animals in a minute. Bold dashed lines in the center correspond to the medians, while upper and lower dashed lines correspond to the quartiles. n is the number of animals analyzed. *t-*test: mock vs nanoalgosome treatment of young animals ns p=0.3387; mock vs nanoalgosome treatment of old animals * p<0.0362.

To validate the nanoalgosome antioxidant effect *in vivo*, we evaluated the response of treated *C. elegans* animals to exogenous oxidative stress. Acute treatment of *C. elegans* with H_2_O_2_ can induce a nearly complete loss of mobility. In fact, 2 hours of treatment with H_2_O_2_ caused a strong decrease in movement (Figure 8e, mock), in contrast to the untreated animals. Interestingly, nanoalgosome treatment counteracted oxidative stress by increasing movement (Figure 8e). This rescue was similar to one obtained with a positive control, N-AcetylCysteine (NAC) (Figure 8e), a widely used antioxidant agent^58^. *C. elegans* copes with changes in ROS levels by expressing detoxifying genes, such as glutathione S-transferase *gst-4*^59^.

To determine whether nanoalgosomes could also directly affect detoxification gene expression, we used a transgenic strain expressing GFP under the control of the *gst-4* promoter^60^. After chronic treatment with nanoalgosomes, we observed a significant decrease in *gst-4* expression levels, suggesting that nanoalgosomes can modulate detoxification gene expression (Figure 8f and Supporting Figure 10). Moreover, antioxidant molecules are capable of counteracting aging and motility decline^61^. In fact, the animal movement is influenced by aging and starting from three days from hatching (young adult stage) it slowly declines^62^.

Therefore, we decided to assess nanoalgosome antioxidant effects by performing an acute treatment at low concentration (10 μg/mL for 16 h), and we analyzed the locomotion of young adult animals (right after the treatment) and older animals (10 days from hatching, 7 days after treatment). As expected, we observed a physiological decline in locomotion between young and old animals (Figure 8g). Interestingly, nanoalgosomes prevented aging effects by preserving the movement capability in older animals and keeping it similar to the one observed in younger animals (Figure 8g). Taken together, our results indicate that nanoalgosomes are bioactive with a clear antioxidant effect both *in vitro* and *in vivo*, supporting their potential role in counteracting aging at molecular and functional level, having the potential to be used as natural and innovative antioxidant effectors.

### CONCLUSIONS

In conclusion, this study demonstrates that nanoalgosomes are a new type of EVs efficiently uptaken by cells through a clathrin-dependent endocytosis and endowed with unparalleled biocompatibility and stability in living organism, unique tropism and bioactivities, including attributes making them suitable as innate anti-inflammatory and antioxidant effectors. Indeed, nanoalgosomes are immune-tolerated *in vivo* and *ex vivo* and exhibit an anti-inflammatory activity *in vitro.* In addition, nanoalgosomes reduced the levels of reactive oxygen species and prevented oxidative stress in tumoral and normal cell lines and in whole organisms, where they counteract aging, thus suggesting that they are enriched with antioxidant compounds. Furthermore, they come with the capacity to be produced at mass scale, with a renewable and sustainable bioprocess from an edible source. This places this technology in an ideal situation to tap into the enormous and still underexploited potential of EVs as novel biological therapeutics, and eventually for further bioengineering to act as delivery vehicles for therapeutic agents.

## METHODS

Nanoalgosomes are isolated and characterized as previously described in Adamo G. et al., 2021^23^, and the relative methods are reported in Supporting methods.

### CELL LINES

The following cell lines were used for the gene expression, bioactivity, and intracellular trafficking analyses: (i) 1–7 HB2, a normal mammary epithelial and (ii) MDA-MB 231, an epithelial human breast cancer and (iii) THP-1 human monocytic leukemia cell lines (ECACC 88081201). All cells were maintained at 37°C in a humidified atmosphere (5% CO_2_)^23,25^.

### *C. ELEGANS* MAINTENANCE AND STRAINS

Nematodes have been grown and handled following standard procedures under uncrowded conditions on nematode growth medium (NGM) agar plates seeded with *Escherichia coli* strain OP50^63^. Strains used in this work have been provided by the *Caenorhabditis* Genetics Center (CGC): wild-type strain N2 (Bristol variety); EG1285 *oxIs12* X *[unc-47p::GFP; lin-15(+)]*(expression of the Green Fluorescent Protein, GFP, in GABA neurons); VC2405 *chc-1(ok2369) III/hT2 [bli-4(e937) let-?(q782) qIs48]* (I;III) (knockout mutant of *chc-1*); RT1315 *unc-119(ed3) III, pwIs503 [vha-6p::mans::GFP + Cbr-unc-119(+)]* (expression of an intestine-specific GFP marker for the Golgi apparatus); RT476 *unc-119(ed3) III; pwIs170 [vha6p::GFP::rab-7 + Cbr-unc-119(+)]* (expression of an intestine-specific GFP marker for late endosomes); CL2166 *dvIs19 III [(pAF15) gst-4p::GFP::NLS]* (expression of an oxidative stress-responsive GFP). Strain QH3803 *vdEx263 [mec-4p::mCherry; odr-1p::dsRed]* expressing mCherry in mechanosensory neurons was kindly provided by M.A. Hilliard (QBI, University of Queensland, Australia)^64^. BY250 *vtIs7 [dat-1p::GFP]* expressing GFP in dopaminergic neurons was kindly provided by M. Aschner (Albert Einstein College of Medicine, NY, USA)^65^.

### NANOALGOSOME STAINING

Nanoalgosomes were labeled with lipophilic dyes: Di-8-ANEPPS (Sigma-Aldrich) for endocytosis inhibition study and *in vivo* studies in *C. elegans*; PKH26 (Sigma-Aldrich) for intracellular trafficking studies; DiR (Sigma-Aldrich) for *in vivo* studies in mice.

Di-8-ANEPPS-, PKH-26- and DiR-nanoalgosome staining were performed as described^23,25^.

Briefly, for *in vivo* studies in mice, DiR-nanolgosome labeling was carried out by 30 minutes of incubation at room temperature with 3.5 µM of DiR diluted in PBS without calcium and magnesium. The same amount of DiR diluted in PBS was used as negative control of free dye. Next, both samples were analyzed by nanoparticle tracking analysis, to check nanoparticle size and distribution, following the removal of free dye by ultracentrifugation (118,000xg for 70 min at 4°C, twice)^23^.The presence of DiR-fluorescence in nanoalgosome-labeled samples was verified by Odyssey scanner IR-measurements (Supporting Figure 6).

### ENDOCYTOSIS INHIBITION STUDY IN CELLULAR MODELS

To study the specific endocytic mechanism involved during nanoalgosome uptake, 1-7 HB2 cells (5×10^3^/wells) were seeded in a 96-well plate. After 24 h, cells were pre-incubated with pharmacological/chemical inhibitors before nanoalgosome addition. Next, the dose- and time-dependent effect of each inhibitor on nanoalgosome cellular uptake was determined. In particular, cells were pre-incubated with the following inhibitors: dynasore (a clathrin-mediated endocytosis inhibitor) at 10, 30, 60, 80 µM; EIPA (5-[N-ethyl-N-isopropyl] amiloride, a pinocytosis/macropinocytosis inhibitor) at 1, 5, 10, 25 µM; nystatin (a caveola/lipid raft-mediated endocytosis inhibitor) at 5, 10, 25, 50 µM. Cells were pre-incubated with dynasore and EIPA for 60 min at 37°C and washed prior to nanoalgosome addition. Cells were pre-incubated with nystatin for 30 min at 37°C. Next, 10 µg/mL Di-8-ANEPPS-labeled nanoalgosomes were added to cells in the presence of these blocking agents, and were subsequently kept at 37°C up to 3 h. The cells treated with nanoalgosomes without inhibitor treatment were used as negative control, while positive control cells were incubated at 4°C to inhibit all energy-dependent mechanisms. Intracellular fluorescence was monitored after 2 and 3 h of nanoalgosome incubation by spectrofluorimetric analysis using a microplate reader (Glomax, Promega). As vehicle-control, cells were cultured in the presence of methanol. A cell viability assay was performed after each treatment (Supporting Figure 1), and treatments were performed in triplicate.

### NANOALGOSOME INTRACELLULAR LOCALIZATION *IN VITRO*

The MDA-MB 231 cell line was grown at a density of 5×10^3^ cells/well in 12-well plates containing sterile coverslips in complete medium for 24 h. Cells were incubated with 2 µg of PKH-26-labeled nanoalgosomes at 37°C for 24 h. Cells were then washed twice with PBS, fixed with 3.7% paraformaldehyde for 15 min, and washed twice with PBS. Afterwards, cells were permeabilized with 0.1% Triton 100-X in PBS with Ca^++^ and Mg^++^ for 10 m at room temperature. Next, cells were incubated with 1% bovine serum albumin (Sigma-Aldrich) in PBS (blocking solution) for 30 m at room temperature to block unspecific binding of the antibodies. Cells were incubated in a primary antibody (CD63 for the endosomal system, clone MX-49.129.5, VWR, anti-LAMP1 for lysosomes, clone 1D4B, Sigma-Aldrich, and anti-calnexin for ER, clone NB100-1965, Novus Biologicals) diluted in blocking solution for 1h at 37°C. Cells were then incubated with a secondary antibody (AlexaFluor-488; Thermo Fischer Scientific) and diluted in blocking solution (1:50) for 2 h at room temperature. Coverslips were mounted with a drop of Vectashield Mounting Medium with DAPI (Sigma-Aldrich). Nanoalgosome intracellular localization was monitored by confocal laser scanning microscopy (Olympus FV10i, 1 μm thickness optical section was taken on a total of about 15 sections for each sample).

### *IN VIVO* SUBCELLULAR LOCALIZATION AND INTERNALIZATION MECHANISM IN *C. ELEGANS*

For subcellular localization and to identify the internalization mechanism, transgenic or mutant animals were treated *in solido*. ∼20 synchronized L4 larvae were transferred on freshly prepared NGM plates seeded with heat-killed OP50 bacteria and Di-8-ANEPPS labeled nanoalgosome (20 μg/mL final concentration) as described in Picciotto *et al.,* 2022^25^. After 24 h of treatment in the dark, young adult animals were transferred to clean NGM plates with OP50 bacteria to let the animals crawl for 1 h to remove the excess of dye. Animals were immobilized as above and confocal images were collected with a Leica TCS SP8 AOBS microscope using a 40x objective.

### DNA DAMAGE IN VITRO AND IN VIVO

#### Gene expression analyses in vitro

Quantitative analysis of mRNA expression were performed in 1-7 HB2 cells. Specifically, 5×10^3^ cells were treated with 2 µg/mL of nanoalgosomes for 24h for genotoxicity study, in which *ATR, CHECK1, RAD51* genes were chosen to assess the expression of DNA damage-related genes. RNA extraction, first-strand cDNA synthesis and gene expression by real-time qPCR are reported in detail in Supporting methods. Treatments were performed in triplicate.

#### Genotoxicity assay in C. elegans

For germline apoptosis quantification, animals were treated *in liquido* as described afterward, with nanoalgosome at a final concentration of 20 μg/mL, PBS as mock and doxorubicin 2.4 μM as positive control. After 72 h of treatment animals were stained by incubation with 33 µM SYTO^TM^12 (Invitrogen, S7574) for 1 h and 30 min at room temperature in the dark. Animals were cleaned by crawling in fresh plates for 30. For microscopy analysis animals were immobilized as above and the quantification of apoptosis after treatment was calculated as the average number of apoptotic corpses per gonadal arm.

### BIOCOMPATIBILITY *IN VIVO* ASSAYS IN *C. ELEGANS*

For the assessment of nanoalgosome toxicity *in vivo*, *C. elegans* synchronized animals have been treated with PBS as mock or nanoalgosomes at different concentrations (1, 20, 64 and 128 μg/mL), through *in liquido* culturing in 96-multiwell plates for 72 h^25^. Synchronized eggs were obtained by bleaching and resuspended in M9 buffer (3 g KH_2_PO_4_; 6 g Na_2_HPO_4_; 5 g NaCl; 1 M MgSO_4_; H_2_O to 1 L) with 2x antibiotic/antimycotic solution (Sigma-Aldrich, A5955), 5 ng/mL cholesterol and OP50 bacteria. 60 µL of this mix containing approximately a total of 30 eggs were aliquoted in each well. After 72 h the number of living animals and of animals reaching the adult stage was quantified per each replicate, with a minimum number of replicates corresponding to three. For the growth assay, synchronized animals have been treated *in solido* on NGM plates seeded with heat-killed (45 min at 65°C) OP50 bacteria and nanoalgosomes (20 μg/mL final concentration), or PBS buffer as mock. The dilutions were performed considering a final volume of NGM in the plate of 4 mL^25^. ∼20 adults per plate in triplicate were let lay eggs for 2 h to obtain synchronized eggs. Animal stage was quantified in each plate every 24 h for 4 days, and the percentage of animals at each stage was calculated on the total number of animals present on the plate. For the lifespan assay, synchronized animals have been treated *in solido*, as described above, with nanoalgosome at a final concentration of 20 μg/mL and PBS as mock. Viability was scored each 3 days and animals were transferred every 3 days in fresh plates with nanoalgosome or mock, respectively. To test the nanoalgosome effect on brood size and embryonic vitality, 20 animals were treated *in solido* in triplicate, as described above, until L4 larval stage and then transferred to new plates without any treatment. Then, every 24 h animals were moved to new plates without any treatment for all the fertile period of the animals (4 days) and the number of laid and hatched eggs counted every day^66^. To assess a putative nanoalgosome neurotoxicity, animals expressing fluorescent protein in GABAergic, mechanosensory and dopaminergic neurons were treated *in liquido* in triplicate, as described above. After 72 h of treatment animals were immobilized with 0.01% tetramisole hydrochloride (Sigma-Aldrich, T1512) on 4% agar pads for microscopy analysis and the morphology and the total number of visible fluorescent neurons was quantified in each animal ^64,67,68^. Confocal images were collected with a Leica TCS SP8 AOBS microscope using a 20x objective.

### *IN VIVO* EXPERIMENTS IN MICE

All animal experiments were performed on wild type BALB/c and athymic nude mice, male and female, which were purchased from Janvier Labs (France) at 5 weeks of age. Animals were used in accordance with Cellvax approved standard operation procedures and with all national or local guidelines and regulations. Mice were housed in specific pathogen-free conditions with food and water were provided ad libitum, in compliance with the Federation of European Laboratory Animal Science Association (FELASA) guidelines. Housing conditions entailed a 12:12 light:dark cycle, room temperature at 20±2°C°, and 70% relative humidity. Mice were acclimatized for one week and were regularly handled by personnel for gentling and habituating to the procedures. On the day of the experiment, animals were randomly allocated into 3 different treatment groups. For the biocompatibility study, groups contained 2 male and 2 female BALB/c mice (n1=4); for the biodistribution study, groups consisted of 3 male and 3 female athymic nude mice (n2=6). Each group received a different treatment. The results were presented as mean ± standard deviation (SD). Multiple comparisons between the groups were performed using GraphPad software.

#### Nanoalgosome biocompatibility study in mice

BALB/c mice (n=4) were injected via intravenous injection into the retroorbital sinus (using 1mL insulin syringes with a 25G needle) with nanoalgosome formulations (dose 1: 10µg corresponding to 4×10^10^ nanoalgosomes/200 µL/mouse; dose 2: 50µg corresponding to 2×10^11^ nanoalgosomes/200 µL/mouse and control: v/v of PBS/200 µL/mouse).

Animals were examined clinically daily, including clinical signs, behavior, signs of suffering (cachexia, weakening, and difficulty of moving or feeding, etc.) and nanoalgosome toxicity (hunching, convulsions). Body weight was monitored at day 3 and 6 post IV injection.

#### Biochemical and hematological analysis

For each animal, 100μL of blood (in compliance with the National Centre for the Replacement, Refinement and Reduction of Animals in Research guidelines) was collected from the lateral tail vein. Animals were euthanized after the last sampling point with CO_2_ (slow fill rate and organ harvesting performed after confirmation of death). Blood was collected and stored in K2-EDTA BD-Microtainers (Thermo Fisher Scientific), centrifuged at 4°C, at 3000 g for 5 min and the obtained plasma (25μL) stored at −20° C. The blood parameters analyzed were hematocrit, lymphocyte count, creatinine, blood urea nitrogen (BUN), liver transaminase enzymes.

#### Nanoalgosome biodistribution in the mouse model

IVIS spectrum studies: The biodistribution of the fluorescently labeled nanoalgosomes was determined by *in vivo* near-infrared fluorescence imaging using an IVIS Spectrum scanner (IVIS Spectrum CT; PerkinElmer). Athymic nude mice (n=6) were anesthetized with 5% isoflurane in 100% O2 at 1L/min for induction, and anesthesia was maintained with 1.2 minimum alveolar concentration via face mask while mice were injected into the retroorbital sinus (as above) with the DiR labeled formulations (listed above) and images were acquired at different post-injection time points (3, 6, 24, 48h). The IVIS Spectrum scanner was set to fluorescence imaging mode, time and an emission filter positioned at 650nm. The focus was kept stable using subject high of 1.5 cm whereas the temperature of the chamber was set at 37°C. Image analyses (set with a fixed count scales from 1′000 to 40′000 and acquired with a 0.2 s exposure time) were computed by first defining the regions of interest (ROI; a representative image with the ROIs are shown in figure 6), which were kept consistent across images, and then the sum of all counts for all pixels inside the ROI (Total Fluorescence Counts-TFqC; photons/second) was recorded.

### ANTI-INFLAMMATORY ACTIVITY ASSAY

The THP-1 cell line was maintained in culture with RPMI 1640 medium (Gibco Life Technologies, Italy) supplemented with heat inactivated 10% Fetal Bovine Serum (FBS, Gibco Life Technologies, Italy) and 1% antibiotic (penicillin 5,000 U/mL, Streptomycin sulfate 5,000 µg/mL, Gibco Life Technologies, Italy). First, cells were differentiated into macrophages by 72h incubation with 200 nM phorbol 12-myristate 13-acetate (PMA). Next, 1×10^6^ cells/mL THP-1 cells were seeded in 24-well in complete medium. After 24 hours, cells were pre-incubated with nanoalgosomes (0.5 µg/mL) for 4 h. After 4 h of pre-incubation, cells were subjected to an inflammatory stimulus, by adding lipopolysaccharide (10 ng/mL LPS from E. coli O55: B5, Sigma-Aldrich) for 20h. Cells stimulated with LPS for 20h (without pre-incubation with nanoalgosomes) represented the positive controls. In addition, cells were incubated with different amounts of nanoalgosomes (0.5 and 0.1 µg/mL) for 24h. Untreated cells are used as negative controls.

#### Cell viability in vitro

Cell viability was evaluated using the CellTiter 96® AQueous one solution reagent (Promega) according to the manufacturer’s instructions. The mean optical density at 490 nm (OD, absorbance) of each wells was used to calculate the percentage of cell viability relative to negative control. Values were expressed as means±standard deviation.

#### Gene expression analyses in vitro

Quantitative analysis of mRNA expression were performed in THP-1 cells, in which *Interleukin-6* genes were chosen to assess the expression of inflammatory cytokine related gene. RNA extraction, first-strand cDNA synthesis and gene expression by real-time qPCR are reported in detail in Supporting methods. Treatments were performed in triplicate.

#### Inflammatory cytokine production in vitro

Interleukin-6 levels in supernatants were determined using commercially available human IL-6 ELISA kits according to the manufacturer’s protocol (Invitrogen). A detailed description of the procedure are reported in Supporting methods. Experiments were performed in triplicate.

### ANTIOXIDANT BIOACTIVITY STUDY

#### Antioxidant bioactivity assay in cellular models

Intracellular ROS levels of living cells were determined using 2′, 7′- dichlorofluorescein diacetate (DCF-DA; Sigma-Aldrich). DCF-DA is oxidized to fluorescent DCF (2′, 7′- dichlorofluorescein) in the presence of ROS, which can be readily detected by a spectrofluorometer. We performed an antioxidant assay on tumoral (MDA-MB 231) and normal (HB2 1-7) cell lines. Briefly, 4×10^3^ cells were cultured in 96-well microplates for 24 h. Cells were then incubated with different concentrations of nanoalgosomes (0.5, 1 and 2 μg/mL) for 24 h. The medium was removed and cells were exposed to PBS containing 40 μM of DCF-DA and kept in a humidified atmosphere (with 5% CO_2_ at 37°C) for 1 h. Next, cells were treated with/without the oxidative agent H_2_O_2_ (250 μM for 3 h) or TBH (250 μM for 1 h) (tert-butyl hydroperoxide solution, Sigma-Aldrich) in the absence/presence of nanoalgosomes. Untreated cells were used as a control to set the percentage of basal intracellular ROS. After extensive washing steps, fluorescence intensity was quantified using a fluorescence plate reader at an excitation of 485 nm and an emission of 538 nm (GloMax® Discover Microplate Reader, Promega).

The relative percentage of intracellular ROS was normalized with respect to untreated cells (control). A cell viability assay was performed after each treatment (see Supporting methods). For gene expression analysis related to antioxidant activity of nanoalgosomes, quantitative analysis of mRNA expression was performed on 1-7 HB2 cells (as described above) and included *CAT, GPX, GSR, AKR, FTH, ALOX, and NOS* genes. 5×10^3^ cells were incubated with nanoalgosomes (2 μg/mL) for 24 h, then, the medium was removed and cells were exposed to oxidative stress (250 μM of H_2_O_2_, Sigma-Aldrich, Germany) for 3 h. Untreated cells were used as negative control.

#### In vivo antioxidant bioactivity assays in C. elegans

For the assessment of exogenous H_2_O_2_ stress response, animals have been treated *in liquido* as described above for 72 h with nanoalgosome at 20 μg/mL final concentration, PBS as mock and N-AcetylCysteine (NAC) (Sigma-Aldrich, A7250) 2mM as positive control. After treatment, animals were exposed to H_2_O_2_ 5mM and water as control for 2 h, then transferred to fresh NGM plates with OP50 bacteria and the number of moving animals on the total number of animals on the plate was scored.

*gst-4p::GFP* expression was quantified after 72 h of treatment *in liquido*, as described above. 5 animals on each slide were immobilized as above, this time using 100 mM NaN_3_ (Sigma-Aldrich, S8032) to allow the lineup of animals. Epi-fluorescence images were collected with a Leica TCS SP8 AOBS inverted microscope, using a 10x objective and FITC filter. Fluorescence quantification was performed using ImageJ, and the Corrected Total Fluorescence (CTF) was calculated for each image using this formula: (integrated density of the area containing the animals) – [(area containing the animals) x (mean fluorescence of background)]^66^. Representative pictures in Supporting Figure 10 were pseudo-colored using Image J.

The anti-aging effect of nanoalgosome on animal movement was assessed through the thrashing assay on young (3 days from hatching) and old (10 days from hatching). Animals at young adult stage have been treated for 16 h *in liquido* with nanoalgosome at a final concentration of 10 μg/mL and PBS used as mock. After treatment animals were transferred to fresh NGM plates and young animals were immediately analyzed, while old animals were transferred every 2 days in fresh plates until day 10 from hatching. For video recording animals were transferred in 7µL of M9 buffer, left 5 minutes to acclimate and then recorded for 30 seconds. The measurement of thrashing was done counting every change of direction respect to the longitudinal axis of the body^69^. Treatments were performed in triplicate.

## STATISTICS AND REPRODUCIBILITY

Statistical analyses were performed with GraphPad Prism software. The experimental replicates and statistical analyses used for each experiment are described in the figure legends.

## RESOURCE AVAILABILITY

Further information and requests for resources and reagents should be directed to and will be fulfilled by the lead contact, Antonella Bongiovanni.

## ASSOCIATED CONTENT

Supporting information (PDF) contains additional figures, tables and detailed supplementary methods, this file is available free of charge.

## ACKNOWLEDGEMENTS

The Authors thank for *C. elegans* strains M.A. Hilliard (QBI, University of Queensland, Australia), M. Aschner (Albert Einstein College of Medicine, NY, USA), S. Martinelli (ISS, Rome, Italy) and CGC, which is funded by NIH Office of Research Infrastructure Programs (P40 OD010440); for Cryo-TEM images Ingo Lieberwirth (Max Planck Institute for Polymer Research, Mainz, Germany); for histological evaluation Yvan Campos (St. Jude CRH, Memphis, USA).

## Author Contributions

Study concepts and design: GA, PS, SP, ED and AB; microalgae cultures and nanoalgosome isolation and characterization: GA, SP, PG, DPR, AP, ER, SR, VL, PC, CA, NT, RN, KL, SM, PB and MM; nanoalgosome *in vitro* studies and data evaluation: GA, SP, PG, MS, AB; *C. elegans* experiments and data evaluation: PS, GZ and ED; gene expression analyses: AN, SC, PC, VL, NA; mice experiments and data evaluation: MW, GA, IR, AB; manuscript preparation: GA, PS, SP, AB, and ED; approval of the final version of the manuscript submitted: all authors.

