## Supplementary Information for "Harnessing Nature’s nanoSecrets: biocompatibility, biodistribution and bioactivity of extracellular vesicles derived from microalgae"

### - Supporting Figures -

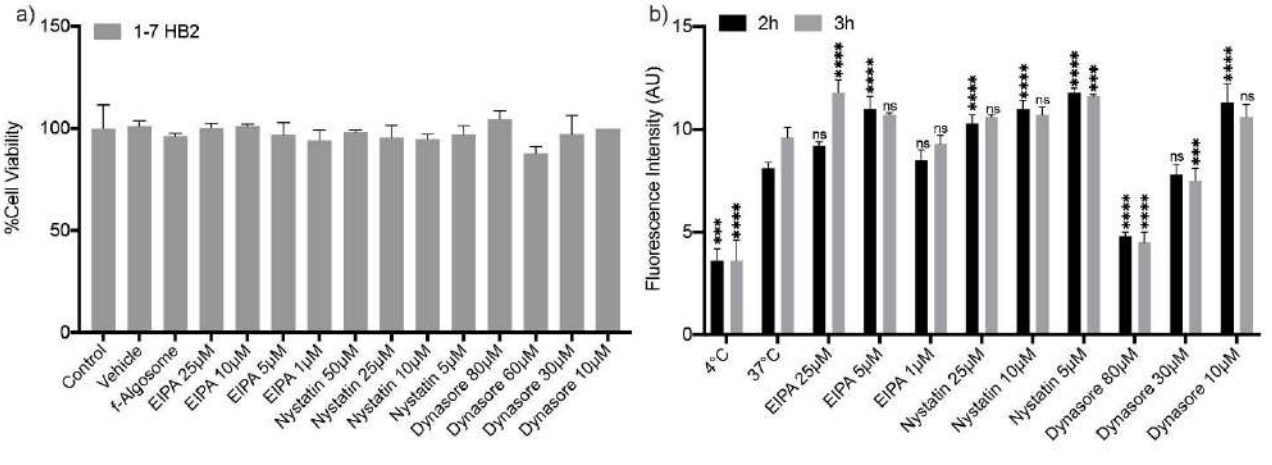


**Supporting Figure 1**. **Cytotoxicity activities and molecular mechanism of nanoalgosome internalization**. **a)** Test of cytotoxicity activities of Di-8-ANEPPS-labeled (f-Algosome) and different concentrations of EIPA (25, 10, 5, 1 µM); Nystatin (50, 25, 10, 5 µM); Dynasore (80, 60, 30, 10 µM) in 1-7 HB2, Cell viability is assessed by using MTS (3-(4,5-dimethylthiazol-2-yl)-5-(3-carboxymethoxyphenyl)-2-(4-sulfophenyl)-2H-tetrazolium) assays. Values were expressed as means ± SD of three independent experiments. By Student's t-test, differences of treated cells were determined not statistically significant when compared with the control. **b)** Effect of metabolic inhibitors of endocytosis on nanoalgosome uptake in 1-7 HB2 cells treated with nystatin, EIPA and dynasore. Results are presented as arbitrary unit of nanoalgosome fluorescence intensity inside cells after 2 and 3h of incubation. One-way ANOVA statistical test was used to assess the statistical significance of the differences: 37°C (2h) (3h) vs 4°C (2h) (3h), EIPA (25,5,1 µM) (2h) (3h), nystatin (25,10,5 µM) (2h) (3h) and dynasore (80,30,10 µM) (2h) (3h), ***p<0,001,****p<0,0001, ns=not significant.

### - Supporting Figures -

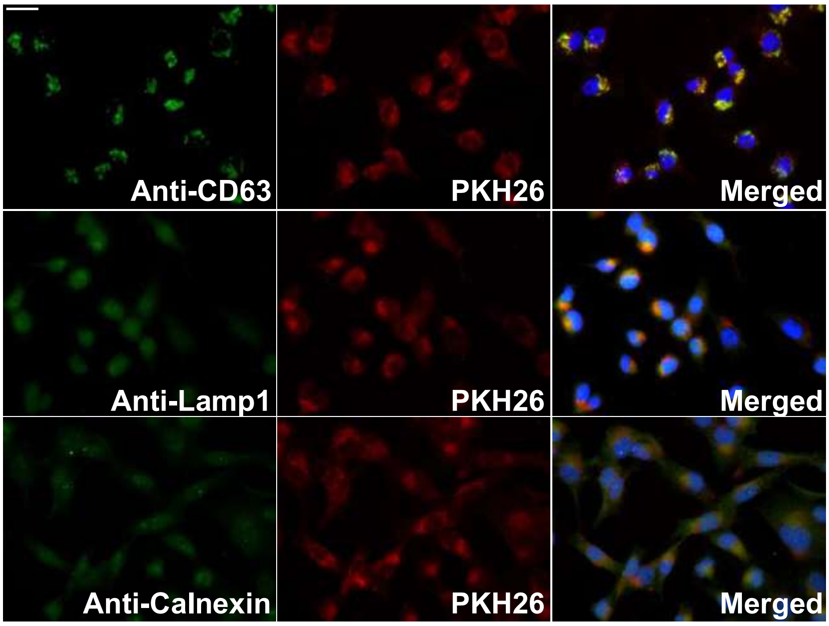


**Supporting Figure 2**. **Intracellular nanoalgosomes localization.** Representative fluorescence microscopy images showing the co-localization of PKH26-labeled nanoalgosome (PKH26) with the endosomal protein CD63 (tetraspanin CD63, green), lysosomal protein Lamp1 (green) and endoplasmic reticulum protein Calnexin (green) in MDA-MB 231 cell lines (nuclei in blue) after 24h of PKH26-nanoalgosome incubation. Magnification 40X. Scale bar 100 µm.

### - Supporting Figures -

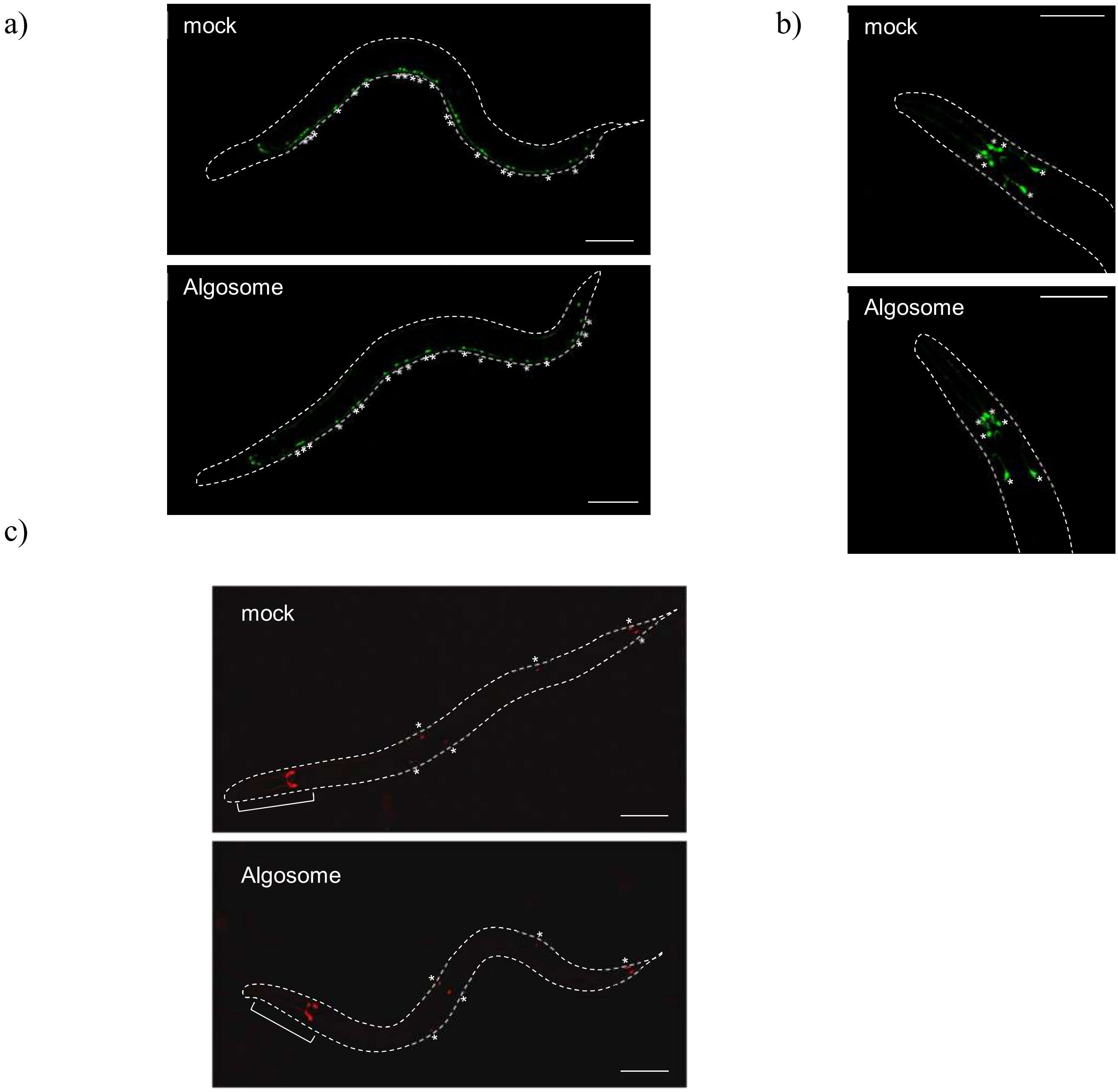


**Supporting Figure 3**. **Nanoalgosome treatment has no effect on neuron morphology and viability**. a) Representative pictures of transgenic animals expressing the GFP in GABA MNs treated with mock (upper panel) or nanoalgosomes (20 µg/mL) (lower panel). Dashed lines mark the shape of the animals, asterisks are in correspondence of each neuron (19 in wild-type). Scale bar is 100µm. Anterior is left, ventral is down.
b) Representative pictures of transgenic animals expressing GFP in the dopaminergic neurons of the head (CEP and ADE) treated with mock (upper panel) or nanoalgosomes (20 µg/mL) (lower panel). Dashed lines mark the head and part of the body of the animal, asterisks are in correspondence of each neuron (6 in wild-type). Scale bar is 75µm. Anterior is left**.** c) Representative pictures of transgenic animals expressing mCherry in the mechanosensory neurons treated with mock (upper panel) or nanoalgosomes (20 µg/mL) (lower panel).

Dashed lines mark the shape of the animals, asterisks are in correspondence of each neuron (6 in wild-type). The injection marker (odr-1p::RFP) is marked by brackets. Scale bar is 100µm. Anterior is left, ventral is down.

### - Supporting Figures –

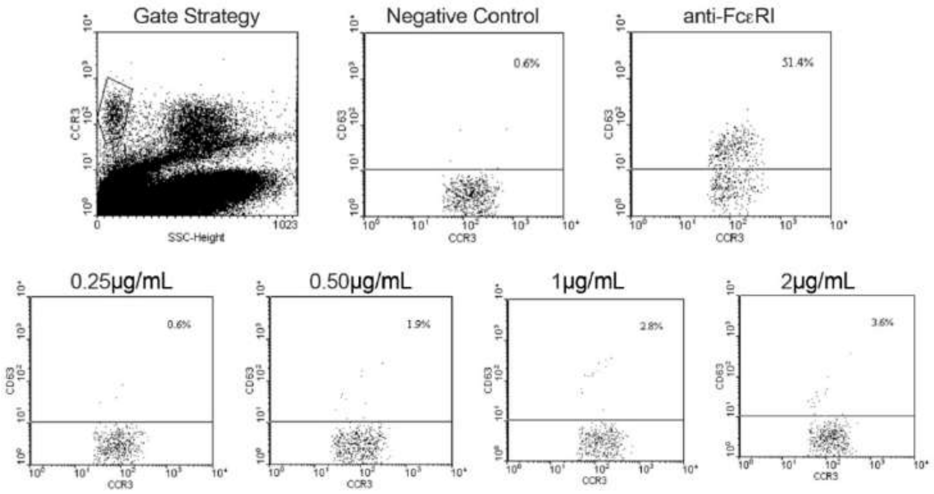


**Supporting Figure 4. Flow cytometry-based basophil activation test.** Upper panels indicate: a representative gate strategy used to select the CCR3 and CD63 populations, and the negative and positive (Anti-FcƐRI) controls. Bottom panels show the activation induced by different concentrations of nanoalgosomes. Representative results of three independent biological replicates are shown.

### - Supporting Figures –

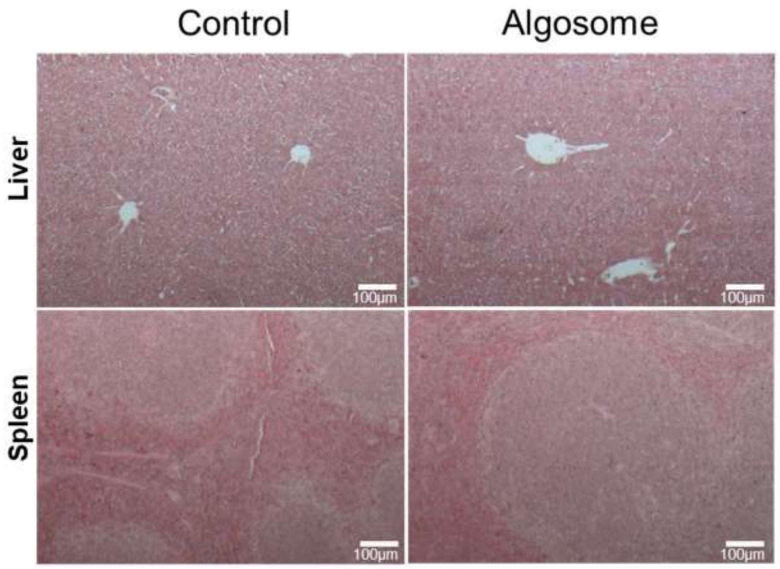


**Supporting Figure 5**. **Nanoalgosome treatment has no apparent histological abnormality.** Representative images of liver and spleen sections from BALB/c mice 48 hours after administration of vehicle (control) or nanoalgosome (10 μg/mouse, corresponding to $4x10^{10}$ EVs/mouse) stained with H&E. No morphological changes in these organs were observed in nanoalgosome-treated mice compared to the control mice.

### - Supporting Figures –

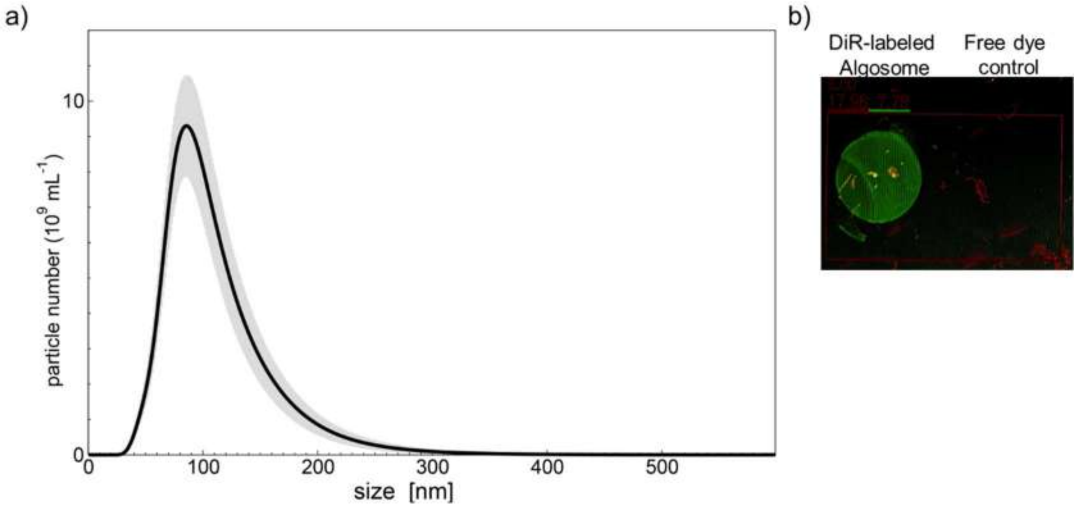


**Supporting Figure 6**. **DiR-labeled nanoalgosomes quality control**. **a)** NTA measurements show a typical size nanoalgosome distribution after the DiR-labeling, with a larger population of vesicles around 100 nm. **b)** The Odyssey IR scanner was used to measure the intensity of infrared fluorescent emissions (λ800nm) of DiR-labeled nanoalgosomes and free dye control (after washing steps to remove free dye).

### - Supporting Figures –

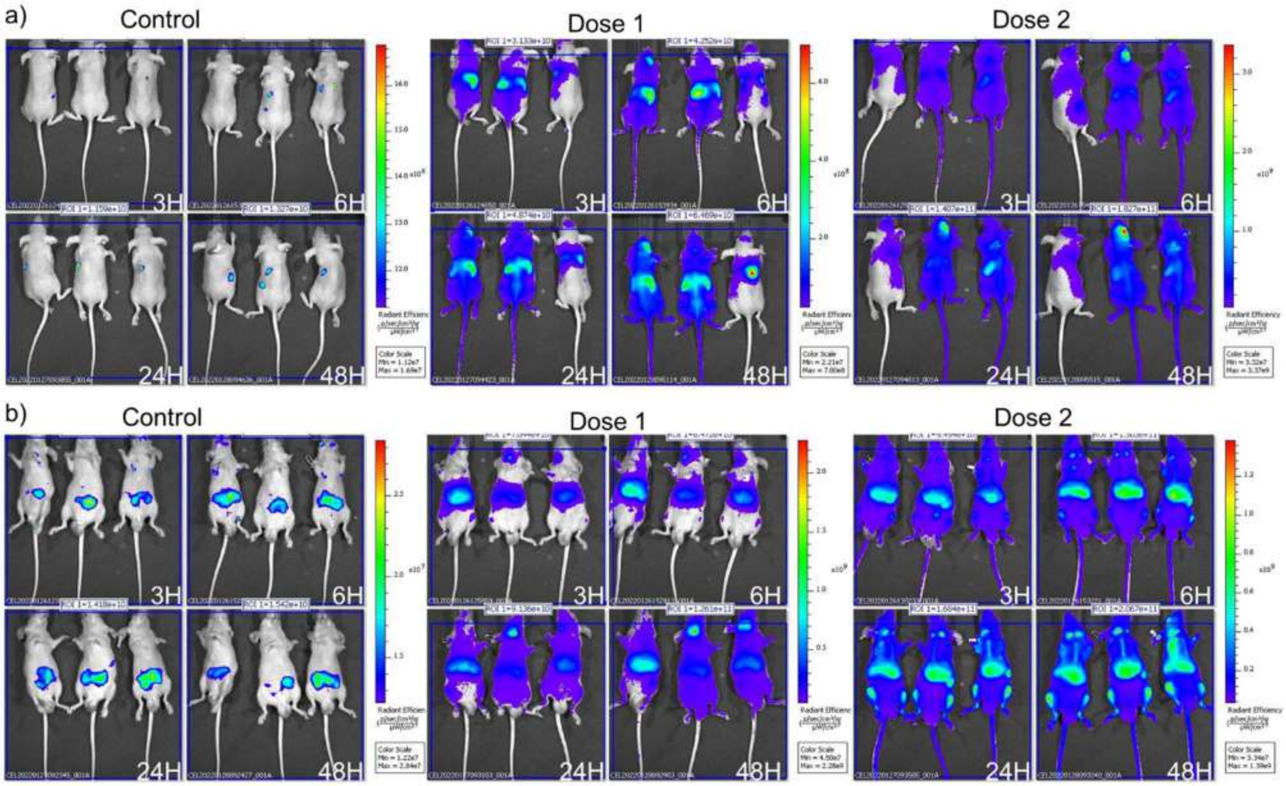


**Supporting Figure 7**. **IVIS imaging of DiR-labeled nanoalgosome biodistribution in nude mice after IV injection**. **a)** Nude mice (n=3; female, prone) and **b)** (n=3; male, supine) nude mice were injected with v/v of free dye control/200 µL/mouse; dose 1: 10 µg/mouse, corresponding to $4x10^{10}$ EVs/mouse; dose 2: 50 µg/mouse, corresponding to $2x10^{11}$ EVs/mouse. Nanoalgosome biodistribution was measured at 3, 6, 24 and 48h post-injection using an IVIS in vivo imaging system at ex/em 644/665.

### - Supporting Figures -

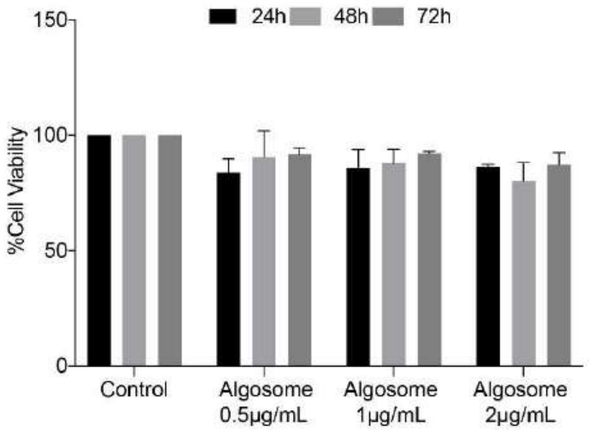


**Supporting Figure 8**. **Cytotoxicity of nanoalgosome in THP-1**. Cytotoxicity of nanoalgosomes in immune-responsive THP-1 cells at different nanoalgosome concentrations (0.5,1, 2 µg/mL) for different time of incubation (24, 48 and 72 h). Values were expressed as means ± SD of three independent experiments. By Student's t-test, differences of treated cells were determined not statistically significant when compared with the control.

### - Supporting Figures –

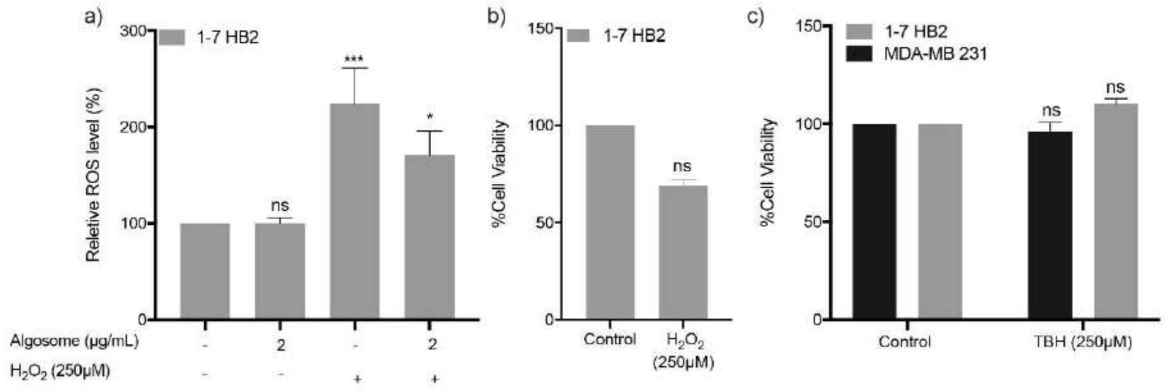


**Supporting Figure 9**. **Antioxidant bioactivity of nanoalgosomes and cytotoxicity activities of oxidant agent in model cellular systems**. **a)** Percentage of ROS production in 1-7 HB2 cells treated with nanoalgosomes (2 μg/mL) for 24h, with/without oxidant agent (250 μM H2O2), normalized to untreated cells. Values are means ± SEM of three independent experiments. One-way ANOVA was used to assess the statistical significance of the differences: 1-7 HB2 vs 1-7 HB2+ 2μgr/mL nanoalgosome, ns=not significant; 1-7 HB2 vs 1-7 HB2 + oxidant agent (250 μM H2O2), ***p<0,001; 1-7 HB2 vs 1-7 HB2 + oxidant agent (250 μM H2O2) + 2μgr/mL nanoalgosome, *p<0.05. **b)** Test of cytotoxicity activities of oxidant agent (250 μM H2O2) in 1-7 HB2 cells and **c)** oxidant agent (250 μM TBH) in -7 HB2 cells and MDA 231. Cell viability is assessed by using MTS (3-(4,5-dimethylthiazol-2-yl)-5-(3-carboxymethoxyphenyl)-2-(4-sulfophenyl)-2H-tetrazolium) assays. Values were expressed as means ± SD of three independent experiments. By Student's t-test, differences of treated cells were determined not statistically significant when compared with the control.

### - Supporting Figures –

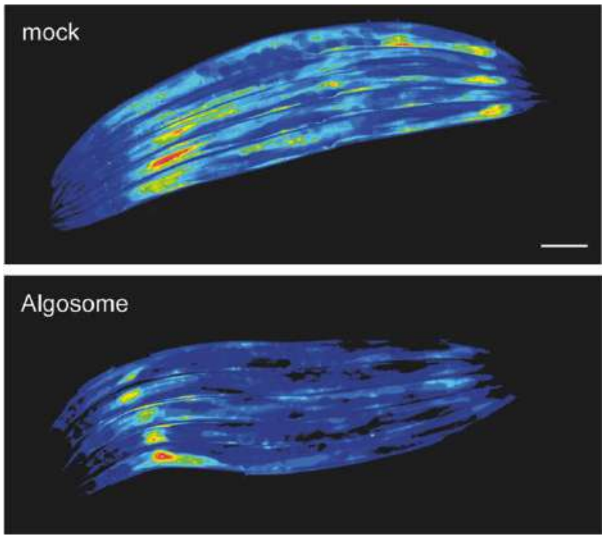


**Supporting Figure 10**. **Chronic treatment with nanoalgosomes decreases the expression level of gst-4**. Representative images of gst-4p::GFP transgenic animals after treatment with nanoalgosomes (20 µg/mL) (lower panel) or mock (upper panel). Images have been false-colored using Image J to allow the visualization of the different fluorescence intensity. Blue colors represent lower signal intensity, and red higher ones. Scale bar is 75µm. Anterior is left.

### - Supporting Tables –

| | Negative Control | Anti-FcƐRI | 0.25µg/mL | 0.5µg/mL | 1µg/mL | 2µg/mL |
|---|---|---|---|---|---|---|
| Subject 1 | 0.6% | 51.4% | 0.6% | 1.9% | 2.8% | 3.6% |
| Subject 2 | 0.3% | 85.9% | 0% | 1.2% | 1% | 2% |
| Subject 3 | 0.9% | 44.2% | 1.6% | 2.3% | 1.5% | 0.9% |

**Supporting Table 1. Flow cytometry basophil activation test.** Table shows the results of basophil test in three subject analyzed. Anti-Fc εRI displays a positive control showing a highly specific monoclonal antibody binding to the high affinity IgE binding receptor. The column labeled 0.25, 0.5, 1, and 2 µgr/mL shows the outcomes following treatment with nanoalgosomes at four distinct concentrations.

### - Supporting Tables –

| | Weight (3 Days) | Weight (6 Days) |
|---|---|---|
| Control | 22,3 | 23,2 |
| | 19,9 | 20,1 |
| | 18,9 | 19,7 |
| | 16,4 | 18,3 |
| | 16,0 | 18,1 |
| | 16,4 | 16,6 |
| **Mean** | **18,3** | **19,3** |
| **SD** | **2,3** | **2,1** |
| Dose 1 | 20,9 | 22,6 |
| | 21,5 | 22,2 |
| | 21,8 | 22,0 |
| | 15,8 | 15,9 |
| | 18,1 | 17,8 |
| | 17,2 | 17,7 |
| **Mean** | **19,2** | **19,7** |
| **SD** | **2,3** | **2,6** |
| Dose 2 | 23,0 | 23,6 |
| | 19,3 | 19,4 |
| | 23,5 | 23,7 |
| | 20,0 | 19,2 |
| | 17,2 | 17,6 |
| | 16,4 | 17,8 |
| **Mean** | **19,9** | **20,2** |
| **SD** | **2,7** | **2,5** |

**Supporting Table 2. Animal body weight.** Mice were weighed every 3 days following nanoalgosome injection.

### - Supporting Tables –

| Gene Name | Primer Forward | Primer Reverse | Gene Bank Accession Number |
|---|---|---|---|
| **AKR1C2** | CAGTGGATCTCTGTGCCACATG | CTGGTTGCAGACAGGCTTGTAC | NM_001135241 |
| **ALOX12** | CCTCGTTATGCTGAAGATGGAGC | ATTTCGGACCCAGGACTTTGCC | NM_000697 |
| **ATR** | GGAGATTTCCTGAGCATGTTCGG | GGCTTCTTTACTCCAGACCAATC | NM_001184 |
| **CAT** | GTGCGGAGATTCAACACTGCCA | CGGCAATGTTCTCACACAGACG | NM_001752 |
| **CHK1** | GTGTCAGAGTCTCCCAGTGGAT | GTTCTGGCTGAGAACTGGAGTAC | NM_001114121 |
| **FTH1** | TGAAGCTGCAGAACCAACGAGG | GCACACTCCATTGCATTCAGCC | NM_002032 |
| **GPX1** | GTGCTCGGCTTCCCGTGCAAC | CTCGAAGAGCATGAAGTTGGGC | NM_000581 |
| **GSR** | TATGTGAGCCGCCTGAATGCCA | CACTGACCTCTATTGTGGGCTTG | NM_000637 |
| **NOS2** | GCTCTACACCTCCAATGTGACC | CTGCCGAGATTTGAGCCTCATG | NM_000625 |
| **RAD51** | TCTCTGGCAGTGATGTCCTGGA | TAAAGGGCGGTGGCACTGTCTA | NM_001164269 |

**Supporting Table 3. The primer sequences used for gene expression in DNA damage and antioxidant studies**.

### - Supporting Methods –

#### Microalgae cultivation and nanoalgosome separation from microalgae-conditioned media

A stock culture of the marine chlorophyte *Tetraselmis chuii* CCAP 66/21b was cultivated in F/2 medium and the purification of nanoalgosomes was carried out following previously described methods.[24]

After a growth period of 30 days, 6 liters of microalgae cultures were clarified using a 450 nm hollow fiber cartridge in the KrosFlo® KR2i TFF System to remove larger particles. The feed flow and transmembrane pressure (TMP) were maintained at constant values of 750 mL/min and 0.05 bar, respectively. The permeate, containing particles smaller than 450 nm, was then subjected to a second filtration step using a 200 nm hollow fiber cartridge, again with a feed flow of 750 mL/min and 0.05 bar TMP. The resulting permeate, containing particles smaller than 200 nm, was subjected to ultrafiltration using a 500 kDa MWCO hollow fiber cartridge, with a feed flow of 750 mL/min and 0.05 bar TMP, and concentrated to a final volume of 150 mL. Subsequently, the samples were concentrated and diafiltered seven times using a smaller 500 kDa cutoff TFF filter module, with a feed flow of 75 mL/min and 0.25 bar TMP, with PBS without calcium and magnesium (Sigma-Aldrich) as the diafiltration solution. This process resulted in a final volume of approximately 5 mL.

#### Nanoalgosome characterization

**BCA assays.** The nanoalgosome protein content was quantified using the micro-bicinchoninic BCA Protein Assay Kit from Thermo Fisher Scientific. This colorimetric method involves comparing the relative concentration of the sample to a protein standard (bovine serum albumin, BSA) and preparing a calibration curve. The relative absorbance of the BCA soluble compound was measured at 562 nm using a GloMax Discover Microplate Reader. The protein content of the nanoalgosomes was determined using the Micro BCA Protein Assay Kit (Thermo Fisher Scientific).

**Nanoparticle tracking analysis (NTA).** The size distribution and concentration of nanoparticles were determined using a NanoSight NS300 (Malvern Panalytical, UK). Nanoalsomes samples were diluted in particle-free water (HPLC-grade Sigma-Aldrich Water filtered by 20 nm using Whatman Anotop filters) to generate a dilution with 20-120 particles per frame. This ensured a concentration within the recommended measurement range of 1-10 × $10^8$ particles/ml. NTA 3.4 Build 3.4.003 (camera level 15-16) was used to analyze five 60-second videos per sample at a syringe speed of 60 in light scattering. Data were further processed using the NanoSight Software NTA 3.1 Build 3.1.46 with a detection threshold of 5.

**Fluorescence nanoparticle tracking analysis (F-NTA).** The Fluorescence nanoparticle tracking analysis (F-NTA) was carried out as describe in Adamo et al., 2020[24]. Briefly, the Di-8-ANEPPS dye was used to label nanoalgosomes, which fluoresces when activated in apolar environments and is specifically enhanced when attached to the lipid membrane of EVs. The fluorescent nanoalgosome (f-algosome) were produced as follows: 5 × $10^{10}$ EV particles/ml were stained with 500 nM of 4-(2-[6-(dioctylamino)-2-naphthalenyl]ethenyl)-1-(3-sulfopropyl)pyridinium, DI-8-ANEPPS (Ex/Em: 467/631 nm, ThermoFisher Scientific), previously filtered by 20 nm filters (Whatmann Anotop filters). Following a 1-hour incubation at room temperature, NTA analyses were performed using a NanoSight NS300 instrument (Malvern Panalytical, UK) equipped with a 500LP filter (laser wavelength 488 nm). The camera level was manually adjusted to optimal settings, and the flow rate for the syringe pump was set at 150 µl/s to ensure that f-algosome crossed the main NTA screen field of view in 5 to 10 seconds. Additional settings were adjusted as described in the previous NTA section. To confirm specificity, a negative control was conducted to verify that the probe alone did not emit a fluorescence signal with F-NTA.

**Atomic force microscopy (AFM).** Atomic Force Microscopy images were performed by using a Nanowizard III scanning probe microscope (JPK instruments) equipped with a 15 µm z-range scanner. Nanoalgosomes were initially concentrated by ultracentrifugation and resuspended in MilliQ water to a final concentration of 5 × $10^{11}$ particles/ml, as estimated by NTA. A 30 µl drop of the sample was incubated on freshly cleaved mica for 10 min, and then gently dried under nitrogen flow. AFM images were acquired in tapping mode by using a NSC-15 (Mikromasch) cantilever (typical spring constant 40 N/m, typical tip radius 8 nm).

**Cryo-transmission electron microscopy (cryo-TEM).** . For imaging, 3 µl of the nanoalgosome preparation ($4 \times 10^{11}$ particles / ml) were applied onto a 400 mesh copper grid covered with quantifoil R2/1 carbon film. Prior to application, the grid was treated with oxygen / hydrogen plasma to enhance hydrophilicity and plunged into liquid ethane using a Vitrobot Mark V (Thermo Fisher). Inspection of the specimen was done using a Titan Krios G4 (Thermo Fisher) transmission electron microscope, operated at 300 kV acceleration voltage. Acquisition was done using a Gatan K3 (Gatan, Pleasonton) camera.

**Multi angle dynamic light scattering (DLS).** An aliquot of vesicle solution was pipetted and centrifuged at 1000×g for 10 min at 4 °C in order to remove any dust particles. The supernatant was withdrawn by pipet tips (previously washed by MilliQ water), put directly into a quartz cuvette and incubated at 20 °C in a thermostated cell compartment of a BI200-SM goniometer (Brookhaven Instruments) equipped with a He-Ne laser (JDS Uniphase 1136P) with wavelength $\lambda = 633$ nm and a single pixel photon counting module (Hamamatsu C11202-050). The autocorrelation function g2(t) of scattered light intensity was measured at a scattering angle $\vartheta = 90°$ by using a BI-9000 correlator (Brookhaven Instruments).

The intensity autocorrelation function g2(t) is related to the size σ of diffusing particles and to their size distribution Pq(σ), by the relation g2(t) = 1 + |β ∫ Pq(σ) exp[−D(σ)q2t]|2, where β is an instrumental parameter, q = 4πñλ−1 sin[ϑ/2] is the scattering vector, with ñ being the refractive index of the medium (ñ=1.3336), and D(σ) is the diffusion coefficient of a particle of hydrodynamic diameter Dh = σ, determined by the Stokes-Einstein relation D(σ) = kBT [3πησ]−1, with T being the temperature, η the medium viscosity, and kB the Boltzmann constant.. The size distribution Pq(σ) is calculated by assuming that the diffusion coefficient distribution is shaped as a Schultz distribution, which is a two-parameter asymmetric distribution, determined by the average diffusion coefficient <D> and its variance V . Two robust parameters may be derived from this analysis: Dz, the z-averaged hydrodynamic diameter (corresponding to the average diffusion coefficient <D>), and PDI, the polydispersity index (PDI = V<D>−2), which is an estimate of the distribution width.

**Immunoblotting analysis.** Proteins of three nanoalgosome batch were separated by sodium dodecyl-sulfate polyacrylamide gel electrophoresis (SDS-PAGE) (10%). A total of 10 μg of nanoalgosome samples (in PBS) were mixed with proper volumes of 5X loading buffer (0.25 M Tris-Cl pH 6.8, 10% SDS, 50% glycerol, 0.25 M dithiothreitol (DTT), 0.25% bromophenol blue). Then, the samples were heated at 100°C for 5 min and loaded in a 10% sodium dodecyl sulfate-polyacrylamide gel for electrophoretic analyses. Proteins were blotted onto polivinilidenfluoro-membranes (PVDF), which were blocked with 3% bovine serum albumin (BSA) in TBS-T solution (50 mM Tris HCl pH 8.0, 150 mM NaCl with 0.05% Tween 20) for 1 h at room temperature. The antibodies anti $H^+$-ATPase (dil. 1:1000 in 3% BSA/TBS-T1x, Agrisera), anti-Alix (clone 3A9, dil. 1:150 in 3% BSA/TBS-T 1X) and anti-Enolase (clone A5, dil. 1:400 in 3%BSA/TBS-T1X), were incubated for 1 h at room temperature. After washing, the membrane was incubated for 1 h with secondary antibodies according to the manufacturer's instructions (horseradish peroxidase-conjugated secondary anti-mouse or anti-rabbit antibodies, Cell Signaling). The membrane were washed four times in TBS-T for 20 min. The immunoblots were revealed using SuperSignal Pierce ECL (Thermo Fisher Scientific).

#### Endocytosis inhibition study

**Cell viability assay.** Tumoral (MDA-MB 231), normal (HB2 1-7) and immune-responsive (THP-1) cell lines were seeded in 96-well plates at a density of $4x10^3$ cells per well and maintained using suitable culture conditions. In particular, for endocytosis inhibition study, we monitored cell viability for 1-7 HB2 cells treated with each inhibitors at different concentrations (EIPA 25,10,5,1 μM; Nystatin 50,25,10,5 μM; Dynasore 80,60,30,10 μM ) as well as with inhibitor vehicle and with nanoalgosomes (f-Alg, 10 μg/ml).

For the antioxidant study, we monitored cell viability in 1-7 HB2 and MDA MB 231 cells treated with H2O2 or TBH at 250 μM. THP-1 cells were treated with different concentration of nanoalgosomes (0.5, 1, 2 μg/ml) for 24, 48 and 72h.

Specifically, 24 hours after seeding, cells were incubated with different concentrations of nanoalgosomes . Cell viability was evaluated using the CellTiter 96® AQueous one solution reagent (Promega) according to the manufacturer's instructions. The mean optical density at 490 nm (OD, absorbance) of each wells was used to calculate the percentage of cell viability relative to negative control. Values were expressed as means±standard deviation of three biological samples, each performed in triplicate.

#### DNA damage *in vitro*

**RNA extraction, first-strand cDNA synthesis and gene expression by real-time qPCR.** Total RNA was extracted from control and treated cells using TRIzol Reagent (Invitrogen), following the manufacturer's instructions, and then stored at −80°C until used. Purified RNA was treated with DNase I, Amplification Grade (Sigma-Aldrich) at 37°C for 30 min to remove any residual genomic DNA contamination, and DNase I was inactivated by adding 50 mM EDTA. First-strand cDNA was synthesized from DNase I-treated RNA samples using the High-Capacity cDNA Reverse Transcription Kit (Thermo Fischer Scientific) following the manufacturer's instructions. The cDNA mixture was tested by PCR using GAPDH and ACTB primers and diluted 1:5 before use in Real-Time qPCR experiments.

**Gene expression by real-time qPCR.** The qPCRs were performed using the BIO-RAD CFX96 system with BlasTaq 2X qPCR MasterMix (Applied Biological Materials) as detection chemistry66. ACTB and GAPDH genes were selected as control genes based on their expression stability in all tested conditions. A normalization factor was calculated based on geometric averaging of ACTB, and GAPDH expression and was used to quantify target gene mRNA levels. Serial dilutions of pooled cDNAs from both control and treated samples were prepared to determine the PCR efficiency of the target and reference genes (data not shown), with amplification efficiency ranging from 1.9 to 2.1. The primer sequences used are listed in Supporting table 3. The qPCRs were conducted according to the manufacturer's recommendations, and each reaction was repeated in triplicate. Amplification conditions were: initial denaturation at 95°C for 3 min and 40 cycles of 95°C for 15 s and 60°C for 60 s, followed by a melting curve from 60°C to 95°C. Amplicons were detected by agarose gel analysis after each PCR to confirm the amplification of the specific gene product (data not shown).

#### Biocompatibility study

***Flow cytometry-based basophil activation test.*** Heparinized peripheral blood was obtained from 3 volunteers. The potential allergenic activity of nanoalgosomes was studied by basophil activation in whole blood samples using flow cytometry to detect the combination of CCR3 and CD63 markers by means of the Flow CAST Basophil Activation Test (Buhlmann Laboratories AG, Switzerland). Blood aliquots (100 µl) were incubated with nanoalgosome dilutions (0.025 - 0.5 - 1 - 2 µg/mL, according to the BCA analysis) in RPMI for 15 minutes at 37°C. PBS and anti-Fc Epsilon RI were used as negative and positive controls, respectively. Results were analyzed using WinMDI 2.8 software (Scripps Research Institute, Jupiter). Basophil activation was reported when more than 5% of activated cells were detected (according to the manufacturer's instructions). The study was approved by the local Ethics Committee (Comitato Etico Palermo 1, 24 February 2021, resolution n. 02/2021).

***Histological examinations.*** For each mouse, spleen and liver were collected, weighed, macroscopic observated, fixed in 4% paraformaldehyde (PFA) and paraffin-embedded. Paraffin sections (thickness of 5-7 µm) sections were deparaffinized in xylene, rehydrated with a graded series of ethanol and then processed for routine hematoxylin and eosin (H&E) used according to the manufacturer's instructions. Finally, all sections were cover slipped using DPX mounting medium and visualized with Nikon Eclipse 80i microscope.

#### Anti-inflammatory Activity Assay

**Cytokine expression - Real Time PCR analysis.** Total RNA was extracted using the QIAzol Lysis Reagent protocol (Qiagen) according to the manufacturer's instructions. The cDNA synthesis was performed using the QuantiTect Reverse Transcription Kit (Qiagen) with 2.5 µg of total RNA. The cDNA was diluted up to 100 µl, and Real Time analyses were performed with an Applied Biosystems StepOnePlusTM Real Time PCR System and Sybr Green technology. Each sample contained 1-100 ng of cDNA in PowerTrack SYBR Green Master Mix (Applied Biosystems, Thermo Fisher Scientific) and 200 nM of specific Quantitect primer assay (Qiagen) in a final volume of 20 µl. The primer sequences used for human cytokines were hIL-6 (Interleukin 6, NM_000600) and the PCR conditions were 95°C for 20 seconds, followed by 40 cycles of two-step PCR denaturation at 95°C for 15 seconds and annealing/extension at 60°C for 30 seconds.

**Inflammatory Cytokine Test.** Medium supernatant and standards were added to the wells and incubated at room temperature for 1h on a horizontal shaker set at 700 rpm ± 100 rpm. Then, the plates were washed and anti-IL-6 conjugate was add into all the wells incubated for 1 h at room temperature on a horizontal shaker set at 700 rpm ± 100 rpm. The plates were washed again and Chromogenic TMB was add to each well. The substrate solution begins to turn blue. After incubation for 15 minutes at room temperature on a horizontal shaker set at 700 rpm ± 100 rpm in the dark, stop solution was added and the optical densities were read at 450 nm wavelength. The results of IL-6 were expressed as concentrations as pg/mL, using standard calibration curve of IL-6. The data were expressed as mean±standard deviation.
